# Scalable Analysis of Multi-Modal Biomedical Data

**DOI:** 10.1101/2020.12.14.422781

**Authors:** Jaclyn Smith, Yao Shi, Michael Benedikt, Milos Nikolic

## Abstract

Targeted diagnosis and treatment options are dependent on insights drawn from multi-modal analysis of large-scale biomedical datasets. Advances in genomics sequencing, image processing, and medical data management have supported data collection and management within medical institutions. These efforts have produced large-scale datasets and have enabled integrative analyses that provide a more thorough look of the impact of a disease on the underlying system. The integration of large-scale biomedical data commonly involves several complex data transformation steps, such as combining datasets to build feature vectors for learning analysis. Thus, scalable data integration solutions play a key role in the future of targeted medicine. Though large-scale data processing frameworks have shown promising performance for many domains, they fail to support scalable processing of complex datatypes. To address these issues and achieve scalable processing of multi-modal biomedical data, we present TraNCE, a framework that automates the difficulties of designing distributed analyses with complex biomedical data types. We outline research and clinical applications for the platform, including data integration support for building feature sets for classification. We show that the system is capable of outperforming the common alternative, based on “flattening” complex data structures, and runs efficiently when alternative approaches are unable to perform at all.

**Key Points:**

- Modern biomedical analyses are integrated pipelines of data access mechanisms and analysis components that operate on and produce datasets in a variety of complex, domain specific formats.
- Scalable data integration and aggregation solutions that support joint inference on such large-scale datasets play a key role advancing biomedical analysis.
- Query compilation techniques that optimize nested data processing are essential for scaling multi-modal, biomedical analysis.

## Background

The affordability of genomic sequencing, the advancement of image processing, and the improvement of medical data management have made the biomedical field an interesting application domain for integrative analyses of complex datasets. Targeted medicine is a response to these advances, aiming to tailor a medical treatment to an individual based on their genetic, lifestyle, and environmental risk factors [21]. The reliability of such targeted treatments is dependent on large-scale, multi-modal cohort-based analyses. This has motivated improved data management and data collection within medical institutions, and has also spurred consortium dataset collection and biobanking efforts [10], which consolidate data sources from hundreds-of-thousands of patients and counting. Examples include 1000 Genomes [3], International Cancer Genome Consortium (ICGC) [24], The Cancer Genome Atlas (TCGA) [55], and UK BioBank [48]. Scalable data integration and aggregation solutions that support joint inference on such large-scale datasets will play a key role in advancing biomedical analysis.

Modern biomedical analyses are pipelines of data access mechanisms and analytical components that operate on and produce datasets in a variety of complex, domain-specific formats. Integrative analyses of complex datasets are difficult to program, especially for large-scale data processing environments. To understand these issues, we now overview multi-omics analysis, distributed processing systems, and the challenges that arise when these two worlds collide.

### Multi-omics analysis

Analyses that combine molecular measurements from multi-omics data provide a more thorough look at the disease at hand, and the relative impacts on the underlying system. For instance, cancer progression can be determined by the accumulation of mutations and other genomic aberrations within a sample [9].

Consider an integrative, multi-omics analysis that aims to identify driver genes in cancer based on mutational effects and the abundance of gene copies in a sample [60]. This analysis combines single-point, somatic mutations and genelevel copy number information to calculate a likelihood score that a candidate gene is a driver within each sample, known as a hybrid score. *Candidate genes* are assigned to mutations based on the proximity of a mutation to a gene. In a naive assignment, candidacy is established if the mutation lies directly on a gene; however, mutations have been shown to form long-range functional connections with genes [42], so candidacy can best be assigned based on a larger flanking region of the genome.

To understand the complexities of such an integrative analysis, first consider the datasources involved. The Genomic Data Commons (GDC) [17] provides public access to clinical information (`Samples`), somatic mutation occurrences (`Occurrences`), and copy number variation (`CopyNumber`). Assume access to each of these data sources returns a collection of objects in JSON (JavaScript Object Notation), a popular format for nested data, where [] denotes a collection type and { } denotes an object type [38].

The `Samples` data source returns metadata associated with cancer samples. A simplified version of the schema contains a sample identifier (`sid`) and a single attribute `tumorsite` that specifies the site of tumor origin; the type of `Samples` is

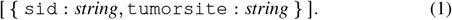

The copy number variation (CNV) data source returns by-gene copy number information for each `sid`; this is the number of copies of a particular gene measured in a sample. The type of copy number information is:

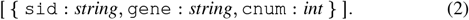

The `Occurrences` data source returns somatic mutations and associated annotation information for each sample. An occurrence represents a single, annotated mutation belonging to a single sample. The type of `Occurrences` is:

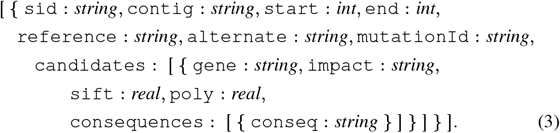

The attribute `candidates` identifies a collection of objects that contain attributes corresponding to the predicted effects a mutation has on a gene; i.e. *variant annotations* sourced from the Variant Effect Predictor (VEP) [33]. The `impact` attribute is a value from 0 to 1 denoting the estimated consequence a mutation has to a gene based on sequence conservation. The `sift` and `poly` attributes provide additional impact scores determined from the Sift [52] and PolyPhen [1] prediction software. These scores estimate the influence a mutation has on functional changes to proteins based on amino acid substitution. The `consequences` foreachcandidategenecontaincategorical assignments of mutation impact based on sequence ontology (SO) terms [13].

VEP provides a distance flag that specifies the upstream and downstream range used to identify gene-based annotations (i.e. the flanking region). This distance flag specifies the size of `candidates`, since more genes are assigned as candidates with a larger flanking regions. A larger value can be used to determine long-range functional connections.

The `Samples` and `CopyNumber` data source types map perfectly into a relational scenario, such as a table in SQL (Structured Query Language) or a DataFrame in Pandas [51]. With all attributes of scalar type (integer, string, etc.), these data sources are considered *flat*. The `Occurrences` data source has a nested collection `candidates` on the first level and another nested collection `consequences` on the second level. When a collection has attributes of collection type, it is referred to as a *complex value* or a *nested collection.*

The hybrid score calculation operates on both flat and nested inputs, associating sample specific copy number values with the impact measurements nested within `Occurrences` based on `gene` and `sid`. After some preprocessing, such an association can be performed with a join in SQL, or a merge in Pandas. The results are then summed to return a collection of candidate genes with corresponding hybrid-scores for each sample. We denote this analysis as `SGHybridScores` with a nested output type:

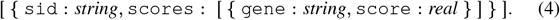

Hybrid scores can be used as risk scores for genes, or these values could be further integrated with additional datasets, such as the interaction of genes in network. The scores can also be further grouped by tumor site, returning sample groups associated to each tumor site. The tumor-grouped hybrid score analysis, denoted as `TGHybridScores`, returns a two-level, nested output with type:

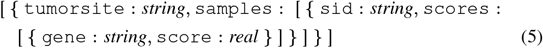

An integrative analysis with nested input and output types, such as the hybrid score analysis, is not straightforward to implement in popular programming languages. Further, this task is exacerbated when the data is large.

### Distributed processing frameworks

Large-scale, distributed data processing platforms such as Apache Spark [59], Apache Flink [4], and Apache Hadoop [12] have become indispensable tools for modern data analysis. The wide adoption of these platforms stems from powerful functional-style APIs (Application Programming Interface) that allow programmers to express complex analytical tasks while abstracting distributed resources and data parallelism. These systems use an underlying *complex data* model that allows for data to be described as a collection whose values may themselves contain collections. Despite natively supporting nested data, distribution strategies often fail to process nested collections at scale, especially for a small amount of toplevel tuples or large inner collections. Further, data scientists who work on local analysis pipelines often have difficulties translating analyses into distributed settings. To better understand these issues, we now provide an overview of a distributed processing framework using Spark as the representative system.

Distributed processing frameworks work on top of a cluster of machines where one is designated as the central, or *coordinator* node, and the other nodes are *workers.* Figure 1 shows the setup of a Spark cluster; an application is submitted to the coordinator node, which then delegates tasks to worker nodes in a highly distributed, parallel fashion. A user never communicates with a worker node directly. A distributed processing API communicates high-level analytical tasks to the coordinator while abstracting data distribution and task delegation from the user.

**Figure 1.**
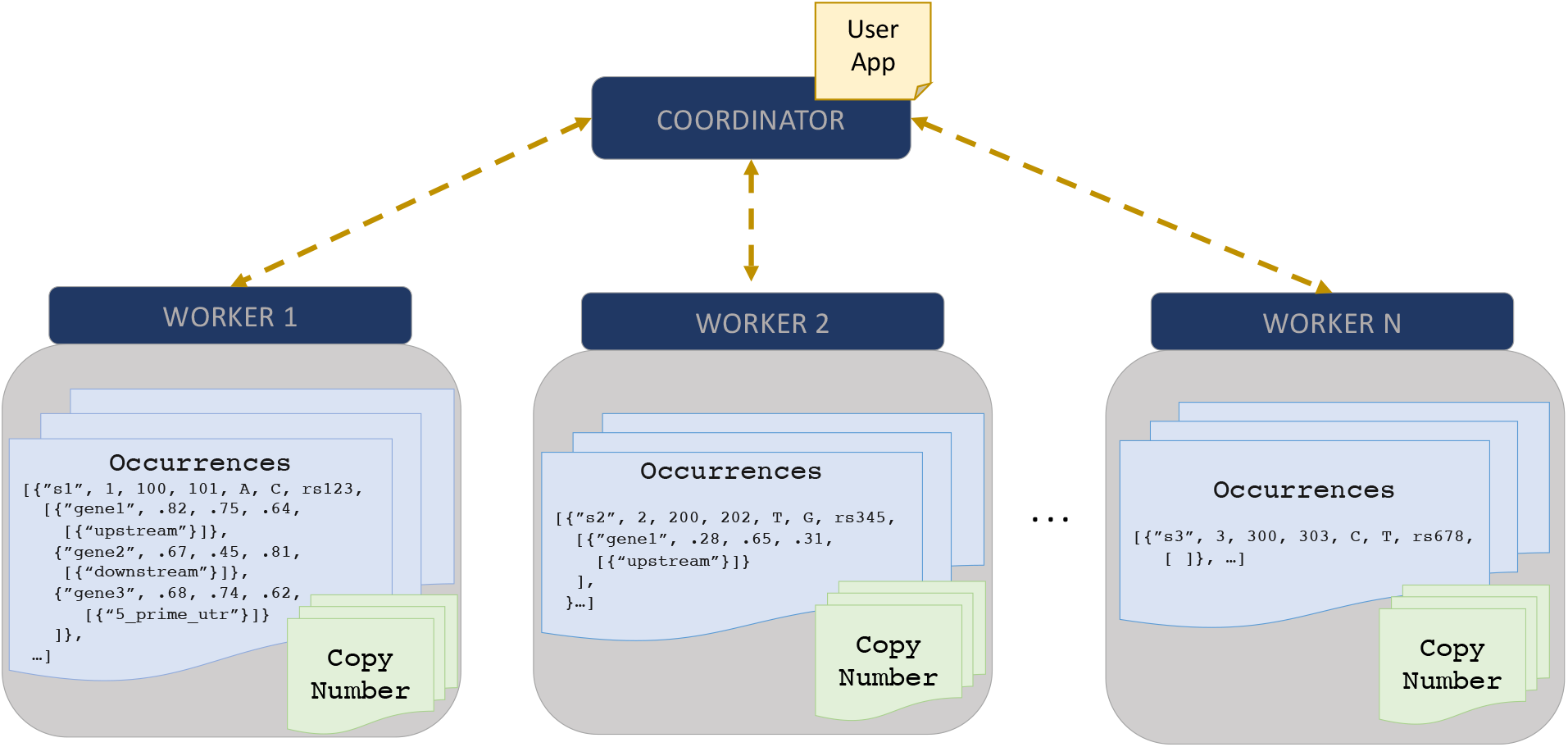
Set up of a Spark cluster with distributed representation of `Occurrences` and `CopyNumber` cachedin memory across N worker nodes. User applications are submitted to the coordinator, which delegates tasks to the worker nodes to support distributed execution. Figure 2 is an example of a user application.

Spark uses a specialized data structure for representing distributed data, known as a Resilient Distributed Dataset (RDD) [58]. An RDD is a collection of distributed objects, where a *partition* is the smallest unit of distribution. When a flat data source, such as `Samples` and `CopyNumber`, is imported into Spark each item of the collection is allocated in round-robin fashion to each partition. The same import strategy is followed for a nested dataset, with the nested attributes persisting in the same partition as their parent. Figure 1 displays how `Occurrences` would be stored in memory across worker nodes, distributing top-level objects with `candidates` and `consequences` nested within the same location. Such a *top-level distribution strategy* ensures all nested values are found within the same partition as their top-level parent.

Spark provides an API for performing batch operations over distributed collections. Figure 2 presents a Scala program that uses the Spark API to associate `CopyNumber` to the relevant gene and samples in `Occurrences`, then groups the result by sample.

The program starts by defining a case class, i.e. a Scala object, denoted `FlatOccurrences` with type [{`sid`: *string,* `gene`: *string*, `impact`: *real* }]. The `flatMap` operation (lines 3-5) works locally at each partition, iterating over top-level objects in `Occurrences` and navigating into `candidates` to create instances of `FlatOccurrence` objects. The join operator (line 6) merges tuples from the resultof `flatMap` and `CopyNumber` based on the equality of (`sid, gene`) values; these are the keys of a *key-based partitioning guarantee* that sends all matching values to the same partition. The process of moving data to preserve a partitioning guarantee is known as *shuffling.* The final `groupByKey` operation (line 7) groups the joined result based on unique `sid` values; this is a key-based operation that sends all tuples with matching `sid` values to the same partition, producing a final output type of: [{ `sid`: *string*, [{`gene`: *string*, `impact`: *real*, `cnum`: *int* }] }].

### Challenges of distributed, multi-omics analyses

Several issues arise when writing programs over distributed, nested collections. First, few top-level values hinder distribution strategies. For example, grouping TCGA-based hybrid scores by the 63 represented tumor sites represented will distribute objects across no more than 63 partitions. This is poor resource utilization for a cluster that supports more partitions. Second, large inner collections, such as candidate genes based on a large flanking region, can overwhelm the physical storage of a partition. This leads to time-consuming processes of moving values in and out of memory. Both of these issues can lead to *skew* related bottlenecks that make certain tasks run considerably longer than others.

Further complications arise when joining on a nested attribute, such as the join between `Occurrences` and `CopyNumber` in the hybrid-score analysis. Since `Occurrences` is distributed, the `gene` join keys are nested within each partition, and thus are not directly accessible without iterating inside the nested collections. Even an iteration inside `candidates` cannot directly perform the conditional join filter on `CopyNumber` because it is itself distributed. An attempt to reference a distributed resource within a transformation of another distributed resource will result in error because a single partition is not aware of the other distributed resources and has no power to delegate tasks to workers. The solution is to replicate `CopyNumber` to each worker node, which can be too expensive, or rewrite to flatten `Occurrences` and bring `gene` attributes to the top-level. Flattening can yield incorrect results; for example, the `flatMap` in Figure 2 will lose all occurrences that have empty `candidates` collections. In general, manual implementations of flattening procedures that ensure correctness are non-trivial [15].

**Figure 2.**
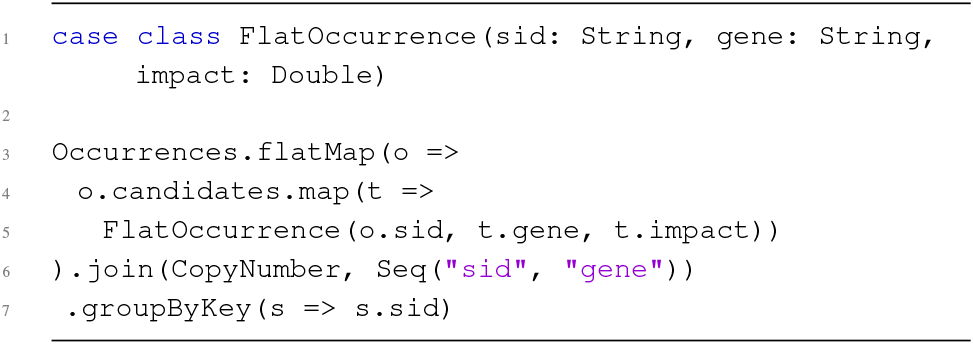
Example Spark application that groups somatic mutations and copy number information by sample.

### Related work

A wide range of tools are available to assist biological analyses. Workflow engines ease the process of connecting many external software systems while producing repeatable analyses; examples include Galaxy [2], Cromwell [53], Arvados [11], and Taverna [37]. Corresponding workflow languages describe imperative pipelines requiring manual optimizations to each individual pipeline component. In contrast, high-level, declarative languages better insulate pipeline writers from platform details, while also providing the ability to leverage database-style query compilation and query optimization techniques. Many genomic-specific languages have been developed that target distributed processing platforms, such as GenoMetric [31], Hail [20], Adam [32, 36], and Glow [18]. These provide advantages for a particular class of transformations, but would not suffice for pipelines that integrate a variety of relational and nested data types.

### Proposed solution

To address these issues and achieve scalable, distributed processing of multi-modal biomedical data, we propose TraNCE (Transforming Nested Collections Efficiently). TraNCE is a compilation framework that automates the difficulties of designing distributed analyses with complex, biomedical data types and provide specific optimizations to ease the difficulty of handling nested collections. The system uses query compilation and optimization techniques and is designed for arbitrary, multi-modal analyses of complex data types.

The paper proceeds as follows. The Methods section outlines the components of the TraNCE platform, describing the major components by means of example. We overview several omics-based use cases that have been trialed with our framework, including performance metrics in the Results section. Finally, we conclude with a summary of contributions and future work.

## Methods

### TraNCE platform

TraNCE is a compilation framework that transforms declarative programs over nested collections into distributed execution plans. This section discusses several key aspects of the platform, including program compilation, program and data shredding, and skew-resilience. *Program compilation* leverages a high-level, declarative source language that allows users to describe programs over nested collections. The framework insulates the user from the difficulties of handling nested collections in distributed environments.

Two compilation routes are provided, standard and shredded, that apply optimizations while transforming input programs into executable code. *Standard compilation* uses unnesting [15] techniques to apply optimal flattening methods in order to compute on nested values. This compilation route automatically handles introducing NULL values and unique identifiers that preserve correctness. *Shredded compilation* optimizes the standard route with *shredding* techniques that transform a program operating on nested collections into a collection of programs that operate over flat collections [7, 56], thus enabling parallelism beyond top-level records.

The result of each compilation route is an Apache Spark program that is suited for distributed execution. We apply dynamic optimizations at runtime that overcome skew-related bottlenecks. *Skew-resilience* prevents the overloading of a partition at anytime during the analysis to avoid such bottlenecks in execution, and maintain better overall distribution of the data.

Figure 3 provides a schematic of the framework, including interaction with a Spark cluster for the shredded compilation method. The Spark cluster setup for the standard compilation method is the classic setup depicted in Figure 1.

**Figure 3.**
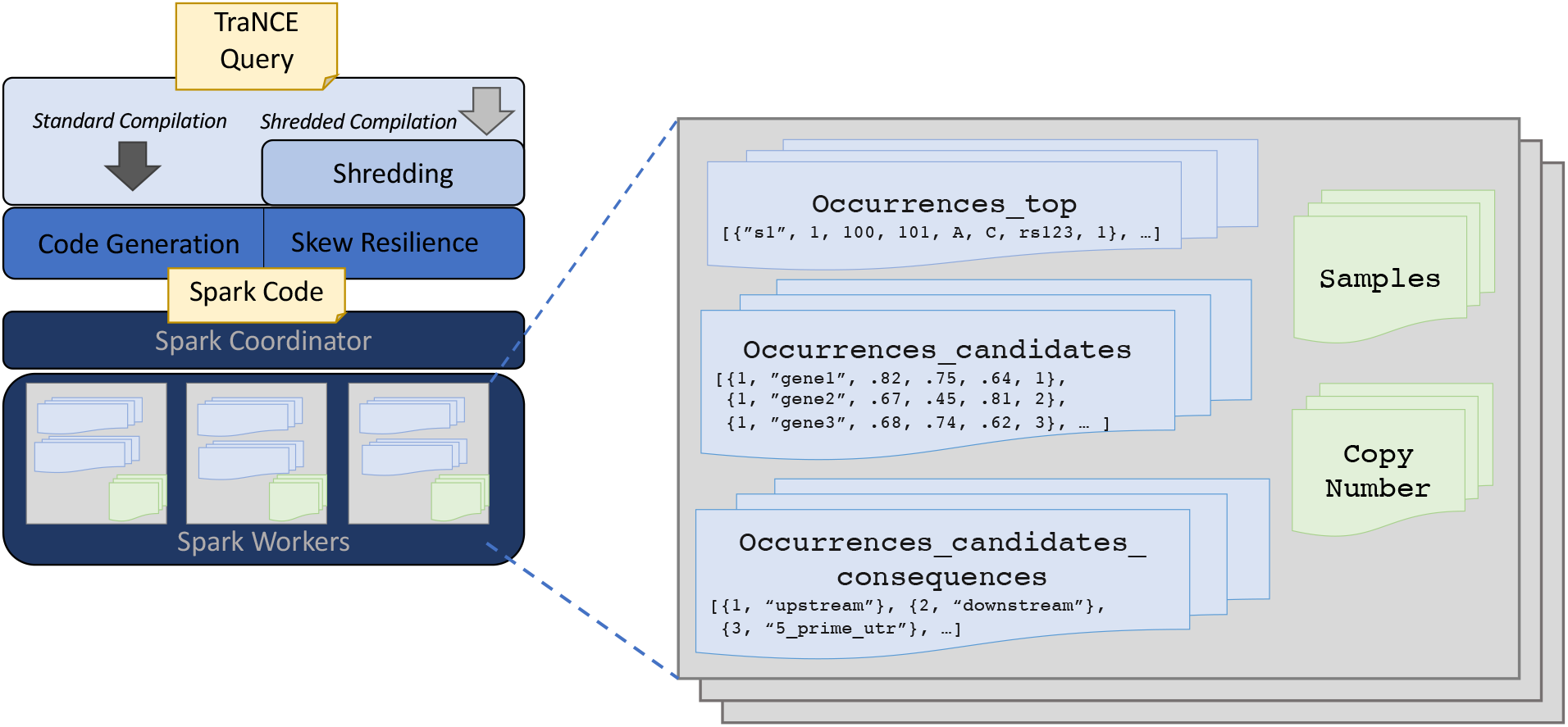
System architecture of TraNCE, presenting two compilation routes that result in executable code. The Spark cluster provides a schematic representation of the shredded compilation route, wherethe shreddedinputs of `Occurrences` are cachedin memory across worker nodes.

The next subsections overview each of the framework components, drawing specific attention to multi-omics analysis. We begin with an introduction to the TraNCE language in the High-level language subsection. The next subsections, Standard compilation and Shredded compilation, review the compilation routes. This section concludes with a note on Skew-resilient processing and Code generation, in there respective subsections.

### High-level Language

For describing biomedical analyses as high-level collection programs, TraNCE provides a language that is a variant of nested relational calculus (NRC) [6, 56]. Here we provide a walk-through of the language using several example programs over the `Occurrences, CopyNumber`, and `Samples` data sources. The full syntax of the TraNCE language is provided in [44].

TraNCE programs operate on collections of objects. Objects are tuples of values for a fixed set of attributes, with all objects of one collection having the same type. Attributes can be of basic scalar type (integer, string, etc.) or of collection type, thus providing support for nested data. We denote collection types with [] and object types with { } to follow JSON syntax. For example, the type of `Occurrences` at (3) is a collection that itself contains collections, with `candidates` and `consequences` corresponding to collection types and all other attributes as scalars.

The main advantage of the TraNCE language is the ability to manipulate nested collections and return results with nested output type, while abstracting out the complications of nested data distribution from the user. Consider the following program, assigned to `OccurrProj` via the ⇐ operator, that requests only specific attributes from the `Occurrences` data source:

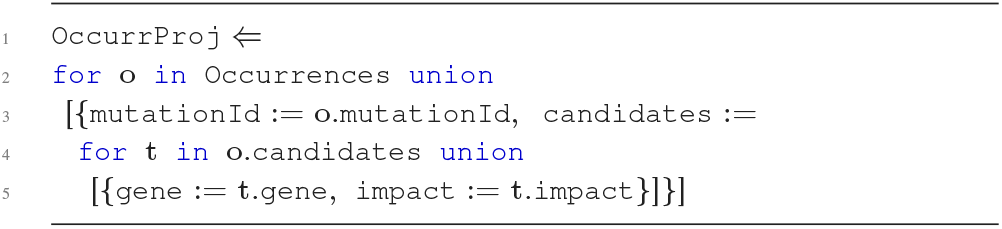

The `OccurrProj` program iterates over the top-level of `Occurrences`, pre-serves the `mutationId` attribute, and creates a nested `candidates` collection by iterating over the `candidates` collection and preserving the `gene` and `impact` attributes.

The language allows one to specify associations between data sources on nested attributes without explicitly defining a flattening operation. For example, the `OccurCNV` program associates copy number information based on both a toplevel attribute and a nested attribute of `Occurrences`.

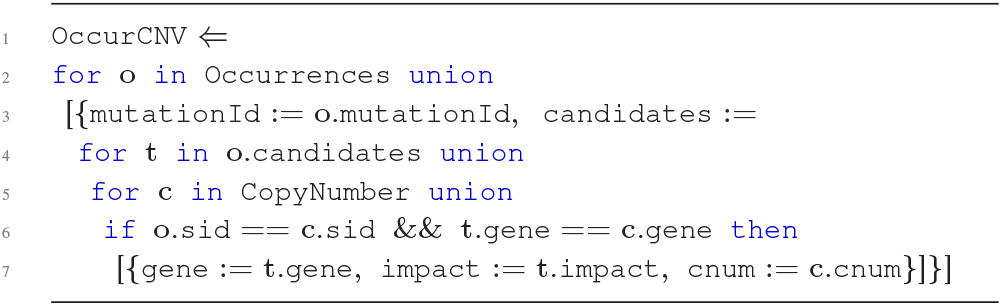

The `OccurCNV` program iterates over `Occurrences` following a structure similar to the previous program. The iteration over `CopyNumber` is specified in the second level, providing immediate access to the nested `gene` attribute and allowing the user to define an association between `sid` and `gene`. The result returns the original structure of the first two levels of `Occurrences`, annotating each of the candidate genes for every mutation with the relevant copy number information.

Standard arithmetic operations and built-in support for aggregation are provided in the language. The 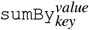 (*e*) function can be used for counting and summing based on a unique key. The *key* and *value* parameters can reference any number of attributes from the input expression *e*. `sumBy` can be applied at a specific level as long as the input *e* has no nesting. For example, the `OccurAgg` program sums the product of copy number variation and mutational impact for every mutation in occurrences.

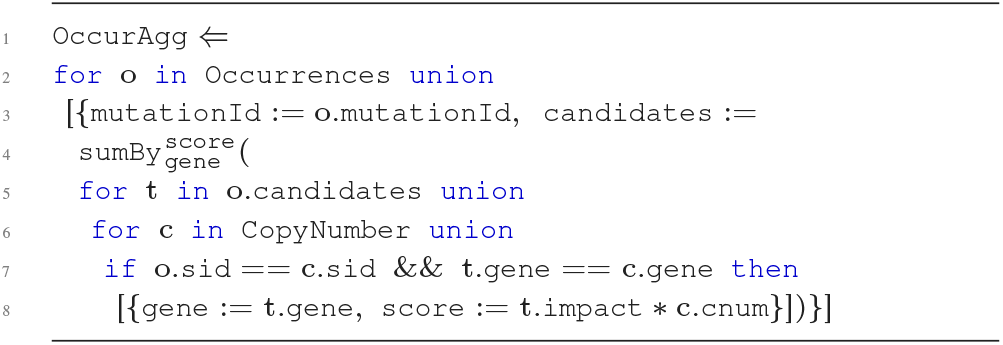

The `OccurAgg` program extends the previous programs, returning the sum of the product of copy number and variant information based on the unique genes in `candidates`. The `sumBy` is applied to the first level of `Occurrences` with `gene` as *key* and `score` as *value*. With all attributes of scalar type, the input corresponding to *e* has a flat type [{ `gene`: *string*, `score`: *real* }].

All programs so far have followed the structure of the `Occurrences` data source, grouping the genes associated to each mutation and each sample. If the goal is to create candidate gene collections per sample, then we can additionally group by sample using the `groupBy`_key_ (*e*) function. The next program `SGHybridScores` will create hybrid-scores for each sample, summing the combination of annotation information and copy number information across all candidate genes for all mutations associated to the top-level sample.

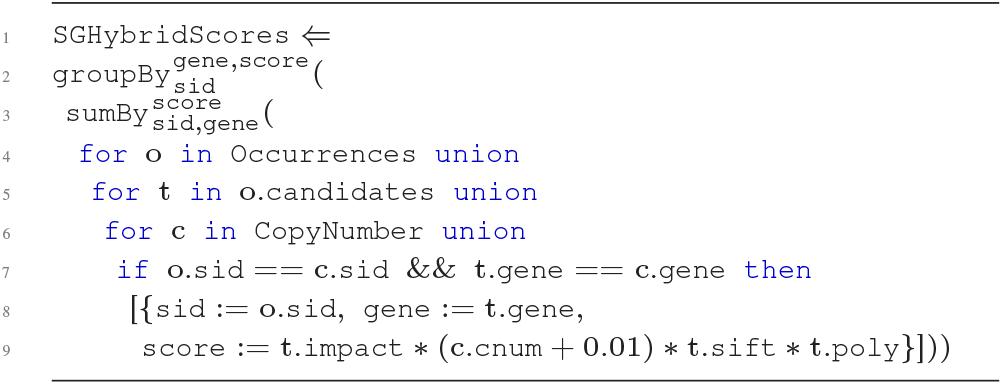

The expression inside of `sumBy` captures the navigation into `candidates`, associating each candidate gene at this level with `CopyNumber` on the `gene` and `sid` attribute. The product of all these measurements produces the intermediate score for each of the candidate genes within the `candidates` collection for each mutation. The final hybrid-scores are calculated by aggregating all intermediate scores across all mutations within a sample, returning a hybrid-score associated to each unique gene for each sample using `sumBy`. The result of this aggregation is further grouped with `groupBy` to return the hybrid scores associated to every sample, producing output type:

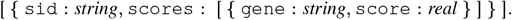

The `Samples` table is used to group the sample-grouped hybrid score results further by tumor site to produce `TGHybridScores`:

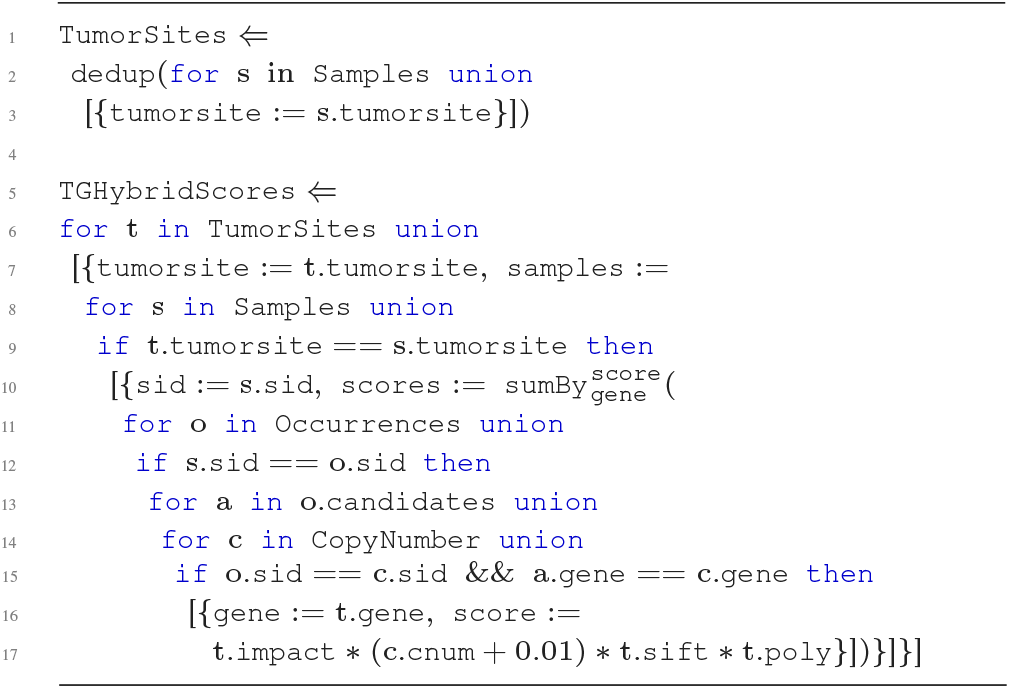

The tumor-grouped hybrid score program first iterates over the `Samples` data source to create a unique set of tumor sites with `dedup. dedup` is a function that returns a collection with all duplicates removed. The second part of the program `TGHybridScores` iterates over these unique groups to create top-level groupings based on `tumorsite`. The program enters the first level at `samples` where it proceeds to iterate over `Samples` creating first-level groupings based on `sid`. The second level begins at `scores` which performs the `sumBy` aggregation described in the previous program. The result of the `TGHybridScores` program is every sample-based hybrid score further grouped by tumor site with the output type:

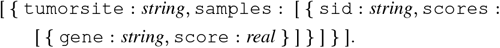

The TraNCE programs described in this section highlight the advantages of using a variation of NRC to describe analyses over nested collections. Each analysis is similar to pseudo-code where the user describes actions on nested collections, without considering implementation details that are specific to a distributed environment.

### Standard compilation

The standard compilation route translates TraNCE programs into executable Spark applications, while handling the difficulties of flattening procedures. Figure 3 provides a high-level schematic of the standard pipeline. This compilation is based on unnesting techniques that automate the flattening process by automatically inserting NULL and unique identifiers (ID) to preserve correctness [15]. The unnesting process starts from the outermost level of a program, recursively defining a Spark execution strategy. A new nesting level is entered when an object contains an expression of collection type. Before entering the new nesting level, a unique ID is assigned to each object at that level. At each level, the process maintains a set of attributes, including the unique IDs, to use as the prefix for the key in grouping and `sumBy` operations.

Consider running the standard compilation for the `TGHybridScores` analysis. The Spark application generated for this program starts by iterating over `Occurrences` values, flattening each of the nested items inside `candidates` with `flatMap`. Prior to this, `Occurrences` is indexed to ensure tracking of top-level objects. If `candidates` is an empty collection, lower level attributes exist as null values. The result of flattening has `gene` attributes that are accessible at top level. The flattened result is joined with `CopyNumber` based on `sid` and `gene` attributes, and the product associated to `score` is calculated. This result is further joined with `Samples` and grouped by sample using the `groupByKey` operation. A final call to `groupByKey` groups again by `tumorsite` to produce the final result.

Projections are pushed throughout the execution strategy, ensuring that only used fields are persisted. The framework can also introduce intermediate aggregations, such as combining `impact, sift`, and `poly` in `Occurrences` prior to joining with `CopyNumber`. The standard route is the basis for our shredded variant and skew-resilient processing module.

### Shredded compilation

The shredded compilation route takes the same high-level TraNCE program as in the standard route, extending compilation to support a more succinct data representation. Analytics pipelines, regardless of final output type, produce intermediate nested collections that can be important in themselves: either for use in multiple followup transformations, or because the pipeline is expanded and modified as data is explored. The shredded pipeline ensures scalability throughout the duration of the pipeline, removing the need to introduce intermediate grouping operations with the help of this succinct representation.

The shredded compilation employs the *shredding transformation*, which transforms programs that operate on nested data into a set of programs that operate on flat data; the resulting set of programs is the *shredded program*. Nested inputs are therefore required to be encoded as a set of flat relations; this is the *shredded input*. The shredded input and shredded program are provided as a succinct representation, where any attribute corresponding to a nested collection is referenced in the flat program using an identifier, known as a *label*. Labels encode necessary information to reassociate the levels of the shredded input. Reassociation is required when a specific level of the shredded program navigates over multiple levels of the shredded input, or when the output is returned as a nested type.

Figure 3 provides a high-level overview of the shredded compilation route, which produces a Spark application that defines the shredded program. The shredding transformation is hidden from the user and a user never interacts with shredded representations directly. Further details of the shredding transformation are described in [44].

Given the transformation to flat representation, the shredded compilation route supports distribution beyond top-level attributes. The succinct representation supports a more light-weight execution that replaces upper and lower-level attributes with labels; this results in reduced data transfer by means of shuffling and provides support for *localized operations,* which are operations that can be directly applied to the level specified in the input program. Shredding can be necessary for scaling for a small number of top-level objects and large/skewed inner collections [44]. Further performance benefits are presented in the Results section. We now continue with an explanation of shredding by example.

The shredded representation of `Occurrences` consists of three data sources:

- a top-level source of `Occurrences`, denoted `Occurrences_top` that returns data with a flat type

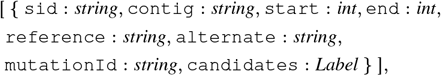
- the first-level source, denoted `Occurrences_candidates`, which has a flat datatype extending the type of `candidates` with a label attribute of *Label* type

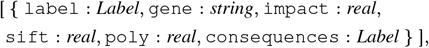
- and the second-level source which extends the type of `consequences` with a label attribute of *Label* type, denoted `Occurrences_candidates_consequences`

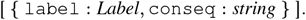

The relationships between the shredded representations can be conceptualized as a database schema, with labels representing foreign-key dependencies. The `candidates` attribute in `Occurrences_top` is then a foreign key that references the primary key of `Occurrences_candidates` at label. Therefore, the reconstruction of nested output, known as *unshredding*, can be achieved by reassociating the shredded sources based on these relationships.

TraNCE then translates the nested program into a series of programs that operate on these flat inputs; i.e. constructs the shredded program. We now review the shredding transformation on the `TGHybridScores` example.

Recall that the tumor-grouped hybrid score analysis starts with the `dedup` operation that returns a collection of distinct tumor sites that will later be used for grouping. The first program returned from the shredding transformation is the shredded program `TumorSites_top`. The expression assigned to `TumorSites` operates over a flat input and returns flat output, so the shredding transformation essentially returns the identity:

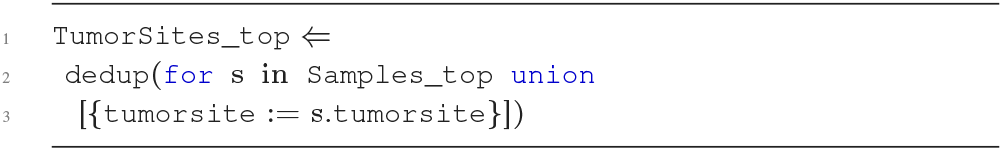

The shredding transformation continues on the expression assigned to `TGHybridScores`, returning a series of three programs; collectively, the shredded `TGHybridScores` program. The first program represents the toplevel collection, `TGHybridScores_top` with the `samples` attribute containing only a label reference.

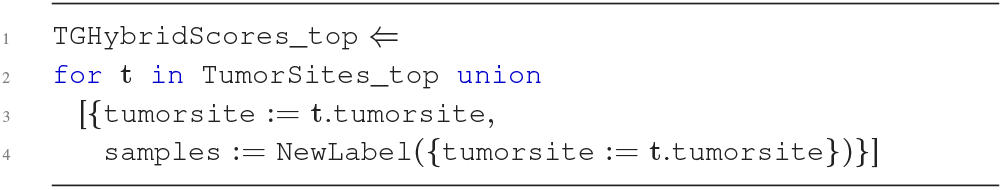

The type of `TGHybridScores_top` is: [{ `tumorsite`: *string,* `samples`: *Label* }]. There are no nested collection attributes, so this is indeed a flat collection. Further, the label of the `samples` attributes encodes only the necessary information to reconstruct the nested output, which in this case is the binding of `tumorsite`.

The program `TGHybridScores_samples` defines the succinct representation of the first-level expression, represented by the following program:

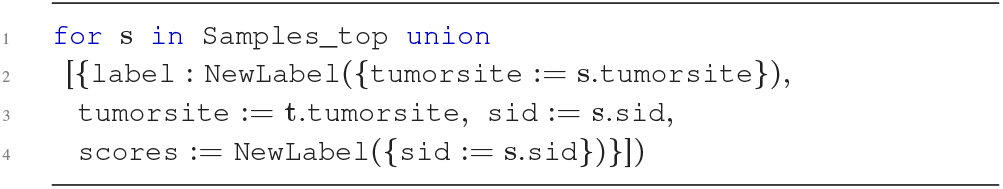

The type of `TGHybridScores_samples` is: [{`label`: *Label*, `sid`: *string*, `scores`: *Label*}].

The `label` expression defines a label that encodes the same information as the `samples` field in `TGHybridScores_top`. This is the same database-style representation seen with the shredded inputs. The label attribute is the primary key of `TGHybridScores_samples` and `TGHybridScores _top` references this with a foreign key at `samples`. The `scores` attribute encodes only the `sid` information that is needed in the next-level expression.

The program `TGHybridScores_samples_scores` defines the succinct representation of the lower-level expression, represented by the final program:

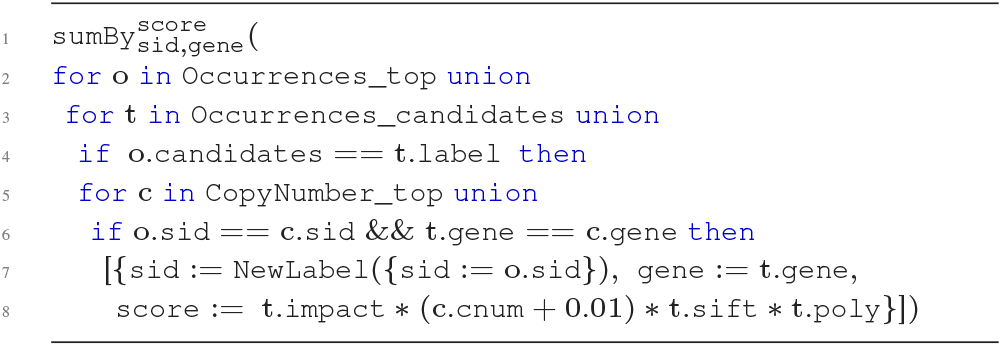

The type of `TGHybridScores_samples_scores` is: [{`label`: *Label,* `gene`: *string*, `score`: *real* }].

The `TGHybridScores_samples_scores` program navigates the top and first level of the shredded representation, using a conditional to reassociate these two shredded representations based on their label attributes. The shredded representation allows nested operations to work directly on the level assigned in the input program. This is an example of a localized operation that results in light-weight execution of nested data, removing the need to carry around redundant data.

The three programs associated to `TGHybridScores` are important for maintaining distribution of the nested values during program evaluation. The first program defines a collection of tumor site information; this avoids distributing the data based on a small number of top-level tuples. The second program defines a collection of sample information, further ensuring the distribution of the lowest level of nesting. The final program defines the bulk of the analysis, isolating the aggregation to the level where it is specified in the input program, and enabling execution of the aggregate without carrying around extra information from the parent.

### Skew-Resilient Processing

Analytical pipelines often contain processes that group items based on a shared attribute, such as the grouping by tumor site in tumor-grouped analysis. The execution of lines 6-7 of `TGHybridScores` will move all data belonging to a specific tumor site to the same node. A TCGA-based `Occurrences` data source will contain significantly more samples for certain tumor sites than others; for example, there are 1100 patients associated to the breast cancer dataset (BRCA) and 51 patients associated to the lymphoma dataset (DLBC). The grouping operation will move all mutations associated to the 1100 BRCA patients to the same node and the whole of DLBC to another node. This will result in extreme imbalances of data across nodes leading to two main issues. First, the movement of a large amount of data to the same location could completely overwhelm the resources on that node - which is likely the case for breast cancer data. Second, any downstream computation of these groups will lead to significant bottlenecks in execution time; a simple count operation over the 444000 BRCA occurrences takes 32x that of the 8115 somatic occurrences of DLBC. Regardless of the specific operation, these distribution issues are a consequence of *skew*. Skew-related issues can easily burden an analysis and can be hard for high-level programmers to diagnose.

Skew is a consequence of key-based partitioning, sending all values with the same key to the same partition. The framework uses a sampling procedure to identify skew-related bottlenecks and automatically handles the distribution of those values at runtime. Given that the shredded representation ensures distribution of inner-collections, the shredded compilation method better handles skew-related issues that arise due to large nested collections and/or top-level distribution. Both pipelines leverage skew-handling methods that maintain proper distribution of values associated to heavy keys. The skew-handling procedure, in general, is beyond the scope of this paper. Further details on the skew-handling methods can be found in [44].

### Code generation

The code generation stage translates a TraNCE program into a parallel data flow described in the Spark collection API, such as the application in Figure 2.

Input and output collections are modeled as Spark *Datasets,* which are strongly-typed, special instances of the RDDs. Datasets are used because the alternative encoding – using RDDs of case classes – incurs much higher memory and processing overheads [43]. Datasets map to relational schemas and also allow users to explicitly state which attributes are used in each operation, providing valuable meta-information to the Spark optimizer.

Since nested inputs are represented as a collection of flat relations in the shredded pipeline, the shredded representation of data sources is merely a collection of Spark Datasets. The tables representing the nested levels contain a label column (`label`) as key; these collections have a label-based partitioning guarantee, which is a key-based partitioning guarantee where all values associated to the same label reside on the same partition. Top-level collections that have not been altered by an operator have no partitioning guarantee and are distributed by the default, round-robin strategy.

The code generator can produce both Spark applications and Apache Zeppelin notebooks. Spark applications generate a single application file that can be executed via command-line. Notebooks can be imported into the Apache Zeppelin web-interface where users are able to further interact with the outputs of the generated code. Notebook generation was designed to provide initial support for users to interface with external libraries, such as pyspark [39], scikit-learn [40], and keras [27]. The notebooks rely on Zeppelin to translate Scala Datasets into Pandas DataFrames for easier interaction with machine learning and other advanced statistical packages. This is merely a first step towards integrating more advanced analytics in the system.

## Results

This section presents a collection of TraNCE programs, coupled with performance metrics, that illustrate the different features of the platform. The first use case is a single-omics analysis that builds mutational burden-based feature sets for use in external learning frameworks. The second use case is a multi-omics analysis pipeline that identifies driver genes in cancer, using nested input and constructing nested intermediate results to return flat output. The first two use cases focus on research applications. The third use case focuses on clinical applications and is designed to mimic requests a clinician could make from a user-interface that supported multi-omics data integration.

The following sections provide details for each use case on the TraNCE platform. We present the performance of each use case using the standard and shredded compilation routes, where standard compilation is used as a representative of flattening methods. Previous results have shown that the standard compilation of TraNCE out performs several external competitors, including SparkSQL [43]. To reflect the reality of compute resources, the first use case is executed with a smaller, single-node environment and the second two use cases on a larger, mutli-node cluster. We now provide an overview of the additional data sources leveraged in these use cases, and proceed with use case description and runtime performance. The final section presents an overview of how the shredded representation can leverage sharing.

### Input data sources

At this point we have presented three data sources used in biomedical analyses: `Occurrences, CopyNumber`, and `Samples`. Here, we describe the additional inputs for each use case and their data types.

#### Variants

The `Variants` data source is based the VariantContext [22] object, used to represent variants from a Variant Call Format (VCF) file. This data structure represents one line, i.e. one variant, from a VCF file. Variants are identified by chromosome, position, reference and alternate alleles, and associated genotype information for every sample. We use an integer-based categorical assignment to genotype calls to support analyses; 0 is homozygous reference with no mutated alleles, 1 is heterozygous with 1 mutated allele, and 2 is homozygous alternate with 2 mutated alleles. The type of `Variants` is:

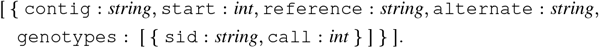

#### Somatic mutations

Somatic mutations are stored in the GDC in Mutation Annotation Format (MAF), which is a flat datadump that includes a line for every mutation across all samples. The type of `Mutations` is:

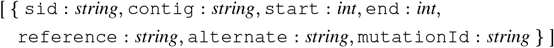

#### Variant Annotations

The variant annotations come from the VEP software, which takes as input mutation information either in VCF or MAF format and returns top-level mutation information augmented with two additional levels of gene and mutational impact information. The overall structure is similar to the `Occurrences` data source (3), except VEP returns a unique set of variant annotations that are not associated to a specific sample. The type of the `Annotations` data source is:

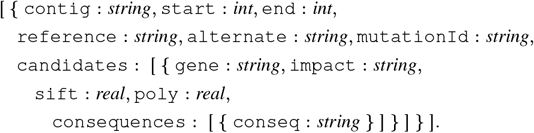

#### Protein-protein interaction network

This `Network` input is derived from the STRING [50] database, which provides a likelihood score of two proteins interacting in a system. The network is represented with a top-level node object and a nested bag of edges. Each edge object contains an edge protein and a set of node-edge relationship measurements. The type of `Network` is:

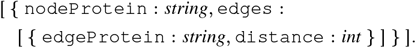

#### Gene expression

Gene expression measurements are derived by comparing transcript counts in an `aliquot` to a reference count. The expression measurement is a normalized count, Fragments Per Kilobase of transcript per Million mapped read (FPKM). The type of `GeneExpression` is:

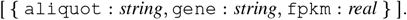

#### Pathway

Pathways are represented as a set of genes. Pathway information is downloaded as a list of curated gene sets from The Molecular Signatures Database (MSigDB) [34, 47]. The type of `Pathway` is:

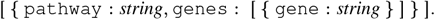

#### Sample Metadata

The `Samples` input maps samples to their aliquots; for the sake of these use cases `sid` maps to a patient and `aliquot` associates each biological sample taken from the patient. Note that this is an extend version of `Samples` introduced at (1). The type of `Samples` is:

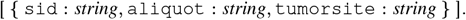

#### Sequence Ontology

The `SOImpact` input is a table derived from the sequence ontology [13] that maps a qualitative consequence to a quantitative consequence score (`conseq`). This is a continuous measurement from 0 to 1, with larger values representing more detrimental consequences. The type of `SOImpact` is:

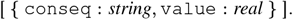

#### Biomart Gene Map

The `Biomart` gene map input is exported from [41]. It is a map from gene identifiers to protein identifiers. This map is required to associate genes from `Occurrences` and `CopyNumber` to proteins that make up `Network`. The type `Biomart` is:

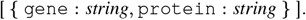

#### Positional Gene Map

Gene mapping files provide the positional location of a gene on a genome, which is a combination of chromosome, start, and end position provided from a General Transfer Format (GTF) file. Each line of the GTF file maps a gene with its positional information [5]. The GTF file can be represented as a flat collection, Genes, with type:

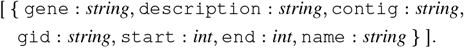

The above data sources are referenced throughout the use cases in the subsections that follow.

### Application 1: Mutational Burden

High mutational burden can be used as a confidence biomarker for cancer therapy [14, 8]. One key measure is *tumor mutational burden* (TMB), the total number of somatic mutations present in a tumor sample. Here we focus on two subcalculations of TMB: gene mutational burden (GMB) and pathway mutational burden (PMB). GMB is the total number of somatic mutations present in a given gene per tumor sample. PMB is the total number of somatic mutations present in a given pathway per tumor sample. These burden-based analyses provide a basic measurement of how impacted a given gene or pathway is with somatic mutations. Mutational burden can be used directly as a likelihood measurement for immunotherapy response [14], or can be used as features for a classification problem.

The progression of some cancers could make it impossible for a clinician to identify the tumor of origin [25]. The ability to classify tumor of origin from a cohort of cancer types can be clinically actionable, providing insights into the diagnosis and type of treatment the patient should receive. For the burden-based use case, we aim to predict tumor of origin from a pancancer dataset.

Figure 4 summarizes the burden-based analyses that calculate GMB and PMB and then perform downstream classification to predict tumor of origin. Each analysis starts by assigning the mutations of each sample, from either `Variants` or `Occurrences`, to the respective gene or pathway. Once assigned, the results are aggregated to return total mutation counts for each gene or pathway producing GMB or PMB values for each sample. The result of the PMB analysis is annotated with tumor site predictor labels from `Samples`, and converted to a Pandas DataFrame to perform two multi-classification methods to predict tumor of origin from a pancancer dataset. We now present the TraNCE programs for these analyses, describe the downstream learning application and discuss performance.

**Figure 4.**
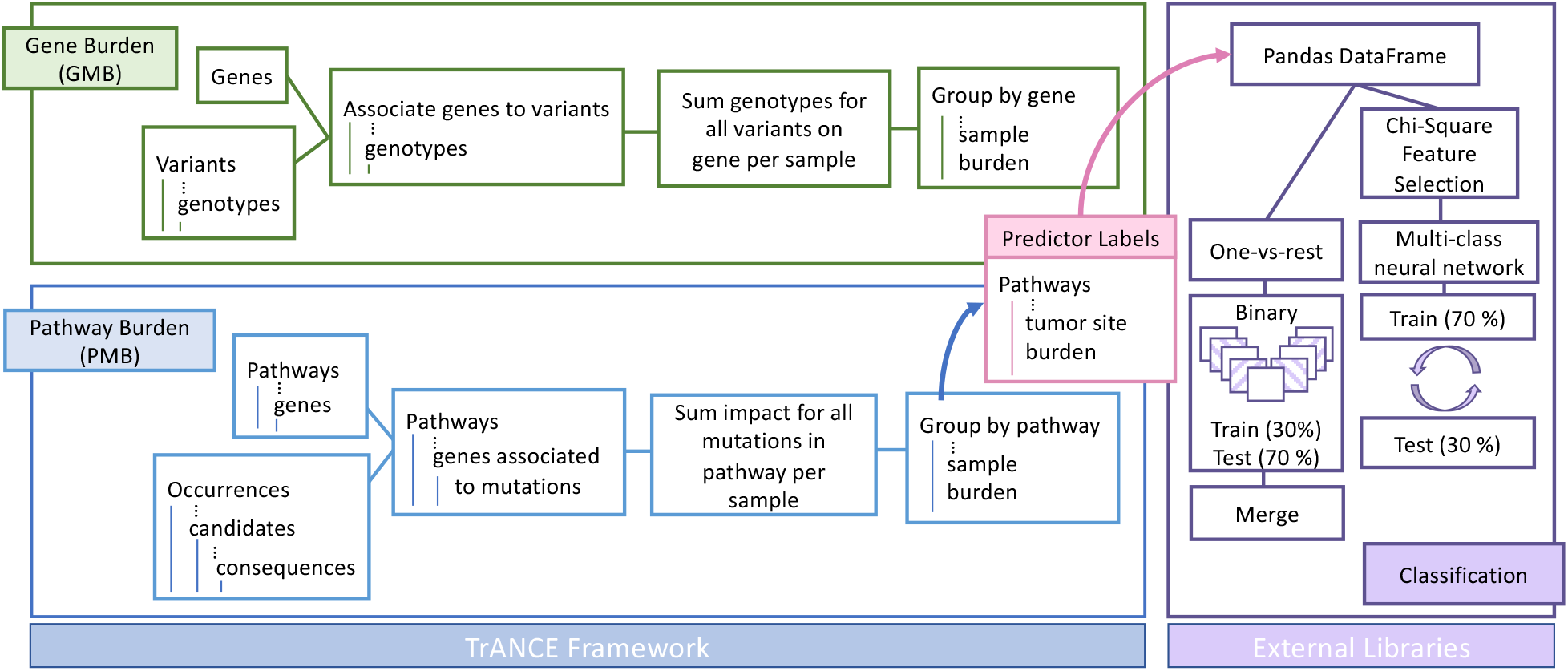
Workflow diagram representing the burden-based analyses for both genes and pathways, and downstream classification problem. The results of the pathway burden analysis feed into a classification analysis using multi-class and one-vs-rest methods to predict tumor of origin.

Given that a pathway is represented as a set of genes, GMB is a partial aggregate of pathway burden; i.e. PMB is the sum of all the gene burdens for each gene in a pathway. We thus show the gene burden program using mutations from `Variants` and the pathway burden program using somatic mutations from `Occurrences`.

#### Gene burden

The gene burden program performs a VCF-based analysis using the `Variants` data source. The program first iterates `Genes` creating a toplevel gene group, and then performs a sum-aggregate of the genotype calls for each sample corresponding to that gene. Variants are associated to a gene if it lies within the mapped position on the genome.

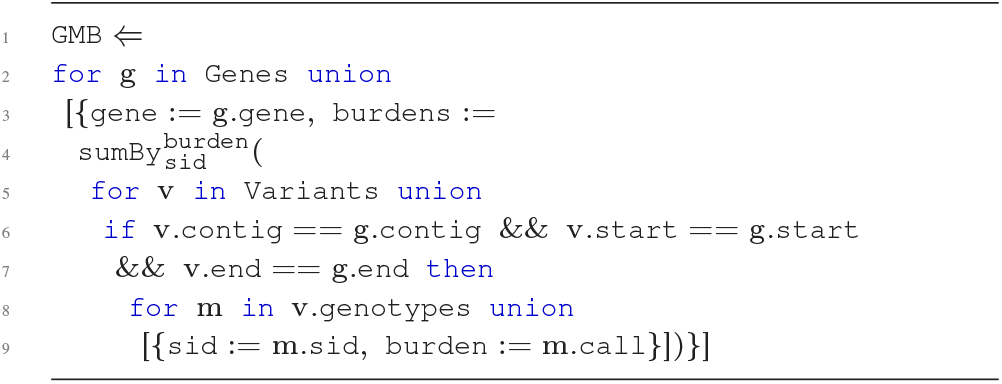

The output type is:

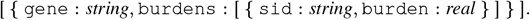

The `GMB` program could be altered to include a larger flanking region by changing the equalities on `start` and `end` to use a range.

#### Pathway burden

The `PMB` program uses the annotations within the `Occurrences` data source to determine gene association. These burden scores are measured within a wider scope than the `GMB` program. When a candidate gene set is created based on a large flanking region, the pathway burdens could be dramatically over-estimated. To account for this, the program uses impact information instead of the number of alleles to measure the mutational burden of a pathway.

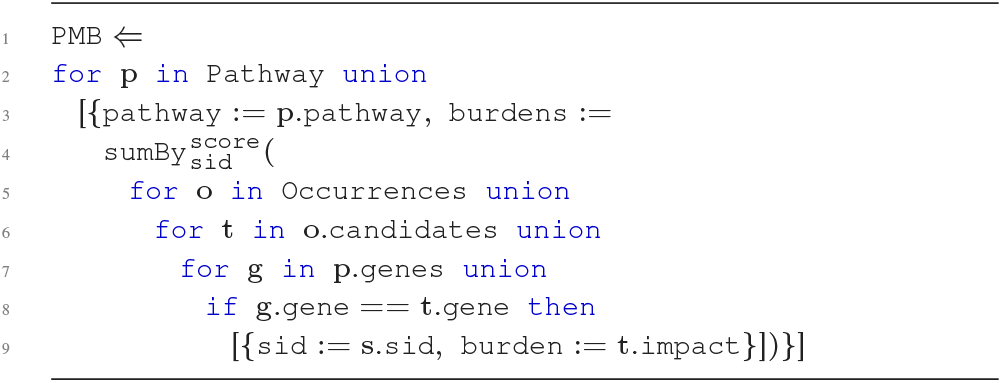

The output type is:

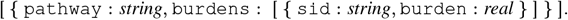

A simple version of the `PMB` program could use raw counts, which we will use for downstream classification analysis. A more complex version could combine multiple impact attributes, such as `impact, poly`, and `sift`, to provide a better estimate of burden.

#### Classification with burden-based features

We now consider how the burden-based programs can be employed to create feature vectors for a learning classifier. Classification of tumor origin has been previously explored with various cancer biomarkers [30, 62, 54, 26]. The goal of our classification problem is to identify tissue of origin from the whole TCGA dataset using pathway burden features based on raw mutation count.

The classification process starts by preparing the PMB output for classification, labeling each pathway burden feature with the associated label:

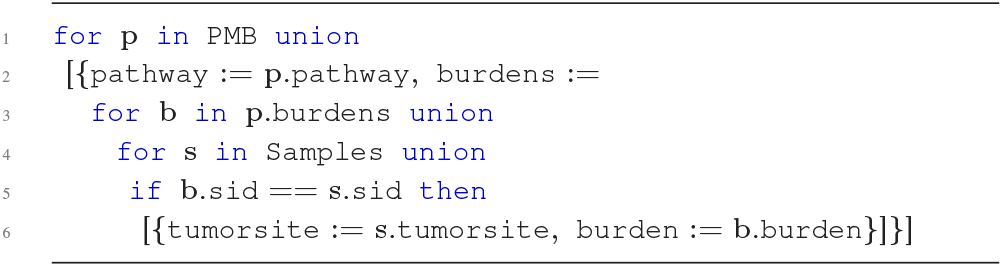

In order to interface with external machine learning libraries, the burden-based programs are compiled into Zeppelin notebooks where the output is available once the program is executed. Learning procedures can then be applied directly in Spark/Scala, or the ZeppelinContext can be used to read a Spark DataFrame as a Pandas DataFrame. For this example, we focus on the Pandas representation to highlight how a user can interact with TraNCE outputs using their external library of choice.

Once represented as a Pandas DataFrame, the data is split for training and testing using scikit-learn and neural networks are constructed with keras. For the whole of the TCGA dataset, we use a minimum cut-off of 200 representative samples. This leaves 9 different tumor tissue sites available for classification: breast, central nervous system, colon, endometrial, head and neck, kidney, lung, ovary, and stomach.

We first train a fully-connected, feed-forward multi-class neural network for tumor tissue site, using 1600 pathways selected by the Chi-squared test as the features. The neural network uses LeakyReLu [57] with *alpha = 0.05* as the activation function, and we utilize dropout layers [46] with *dropouī_raīe = 0.3* after each fully connected layer (dense layer) before the output. This model is trained using the categorical cross-entropy loss function and the Adam optimizer [29]. The network has a Softmax output, which can be interpreted as a probability distribution over 9 different tumor tissue sites. The data is randomly split into two folds, 70% for training and 30% for testing.

Next, we extend the previous method via the “one-vs-rest” method [61], which decomposes a multi-classification problem into multiple binary classification problems and each binary classifier is trained independently. For every sample, only the most “confident” model is selected to make the prediction.

Each binary classifier is a fully connected, feed-forward neural network, using all 2230 pathways as the features. These are set up the same as the multi-class networks, except with dropout layer *dropouī_raīe = 0.15* and a binary crossentropy loss function. The binary networks have Sigmoid output, which can be interpreted as a probability of a certain type of tumor tissue site corresponding to this model. For each model, the data is randomly split into two folds as with the tumor-site network.

We train 9 independent binary classifiers for each type of tumor tissue site. These binary models predict the likelihood that the given pathway burden measurements of a patient are associated with the tumor site represented by that model. After training each binary model, predictions are made using the entire dataset, and the computed results are merged. The probabilities from all models are compared for each patient from the testing dataset, classifying the patient according to the highest likelihood. For example, suppose we have two models, a breast model that predicts a breast-site likelihood of 0.8 and a lung model that predicts a lung-site likelihood of 0.6 for the same patient. The system compares these two probabilities and classifies tumor of origin as breast.

Difference in sampling procedures aside, the multi-classifier and the binary models in the one-vs-rest method have one key difference. When using pathway burden features, pathways that are highly correlated with a specific tumor site could be overpowered by pathways that show strong signal for cancer in general. The multi-classifier could compromise features specific to tumor of origin in an attempt to achieve best performance overall. This can lead to particularly inaccurate results when the data distribution is uneven. The binary models are eager to select the best feature weights for the representative tumor of origin, providing more opportunities for tumor-specific features to stand out.

#### Runtime performance

We perform the burden-based analyses using Spark 2.4.2, Scala 2.12, and Hadoop 2.7 on a machine with one worker, 10 executors, 2 cores and 20 Gigabyte (GB) memory per executor, and 16GB of driver memory. Due to resource limitations of this cluster, we use the publicly available chromosome 22 from phase 3 of the 1000 Genomes Project [3, 49]; this is a 11.2GB dataset representing 2504 samples. The runtime of a program is measured by first caching all inputs in memory.

Figure 5 displays the runtimes of the standard (Standard) and shredded (Shred) compilation for the gene burden and pathway burden analysis for an increasing number of variants from the VCF-based `Variants` data source. The results show that flattening methods of Standard are quickly overwhelmed as the number of variants increase; whereas, Shred increases at a much slower rate. In addition, after 600-thousand variants Standard pathway burden increases at a greater rate than the corresponding gene burden run. The shredded method exhibits two main advantages. First, the succinct representation avoids carrying around extra data, such as the genotypes information when `Variants` are joined with `Genes`. Second, the result of flattening `Variants` will have a large amount of items. The whole file contains roughly 1103600 variants and more than 2500 samples, which produces a result with over 2.7 billion items. These results highlight the advantage of the shredded representation even for the shallow nesting of the VariantContext structure.

**Figure 5.**
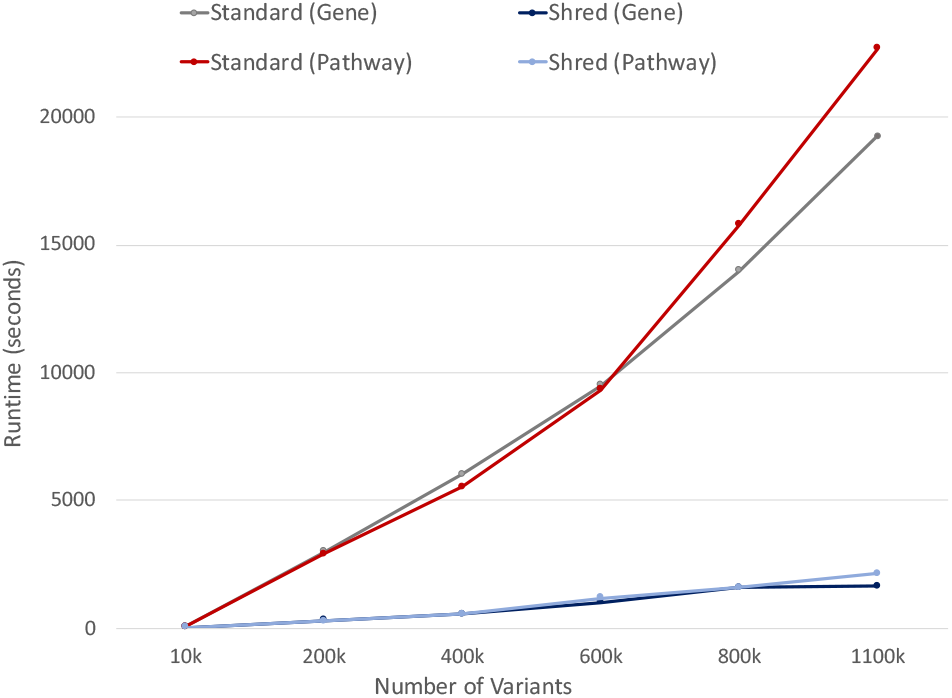
Performance comparison between the standard and shredded pipeline on gene and pathway burden analysis 1000 Genomes dataset.

#### Multi-classification results

Figure 6 shows the accuracy and loss of the multiclass neural network for tumor tissue site for 30 epochs. The overall accuracy is 42.32%, calculated from the confusion matrix adding all 444 correctly predicted labels together and dividing by the 1049 testing samples. Most misclassifications were predicted to be breast cancer, likely attributed to the data imbalance problem of the training dataset. An imbalanced data distribution forces a model to learn features corresponding to highly populated labels, reducing training loss while skewing overall prediction performance.

**Figure 6.**
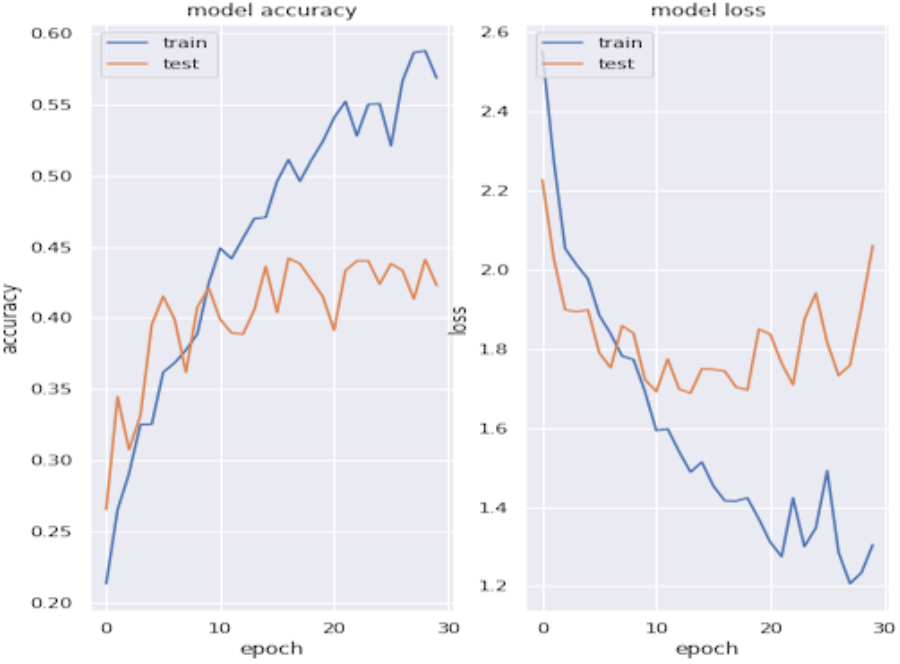
The accuracy and loss of the multi-class neural network for tumor tissue site.

Different types of cancer may not contain enough dominant features for a simple multi-class model to distinguish differences among tumor origin site. Even pathways that play a key role in any cancer, such as pathways specific to disruption in cell-cycle, could not be providing enough signal to act as a determinant for cancer types. This could be because other pathways are washing out the signal of more important pathways, or it could simply mean that pathway burden alone is not providing the whole story. Thus, future multi-class problems in this domain should consider integrating other features, such as additional genomic measurements, or filter pathways based on prior knowledge of the cancer types in question.

#### One -vs-rest classification results

Figure 7 displays the accuracy and loss of three binary networks for 10 epochs. We present the three worst-performing classes from the multi-class network: stomach, head and neck, and central nervous system, which all resulted in testing accuracies above 90% in the one-vs-rest method. The accuracy and loss of the other binary models are provided in the supplementary material. The combined accuracy of all binary models is 78.44%, calculated as the correctly predicted labels (2744) divided by total samples (3498). Overall performance of the one-vs-rest method is far better than the multi-classifier performance.

**Figure 7.**
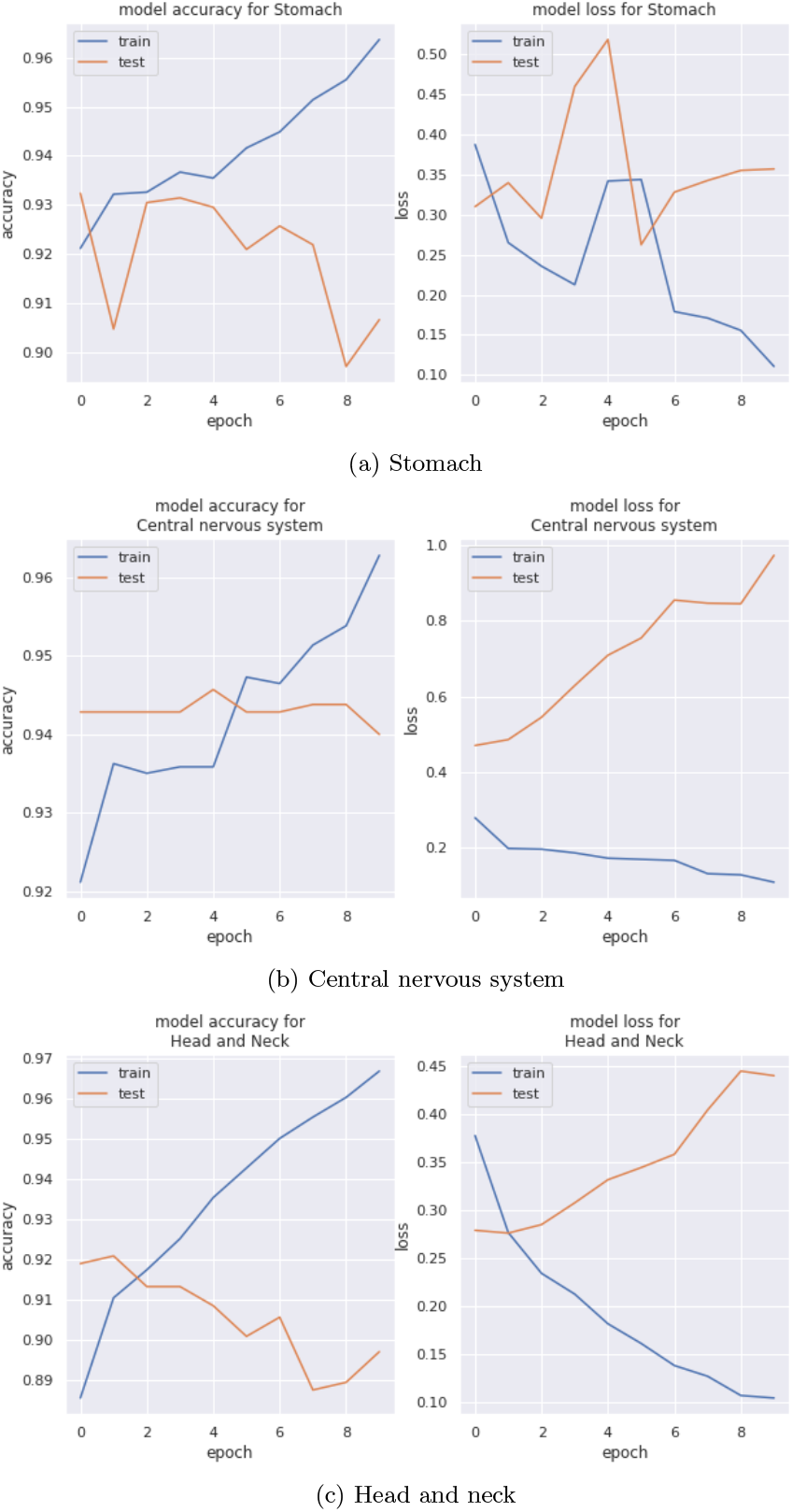
Accuracy and loss for the tumor tissue site based binary network, includes results for the three worst-performing classes from the multi-class network.

Further exploration into pathway signal profiles of each tumor site could be considered for future work. Gene burden performance could be compared to that of pathway burden in order to identify genes that are the main drivers for pathway signal. The identification of predominant pathways and genes for certain tumor sites could provide insight into specific cancer profiles and determine overall confidence of using burden-based features for tumor site classification.

The burden-based use case exemplifies how TraNCE can handle data integration tasks, and more specifically, integration tasks that produce feature vectors for classification problems. In addition, this use case shows how users can interact with popular learning packages within a notebook environment without the overhead associated with manually integrating data sources.

#### Application 2: Multi-omics cancer driver gene analysis

Mutations that play a driving role in cancer often occur at low frequency [19], making cohort analysis across many samples important in their identification. Further, a cancer profile is more than just a consequence of a single mutation on a single gene. Gene interactions, the number of such genes, and their expression levels can provide a more thorough look at cancer progression [9]. This use case focuses on such a multi-omics analysis, which defines a set of programs that integrate annotated somatic mutation information (`Occurrences`), copy number variation (`CopyNumber`), protein-protein network (`Network`), and gene expression (`GeneExpression`) data to identify driver genes in cancer [60]. This analysis provides an integrated look at the impact cancer has on the underlying biological system and takes into account the effects a mutation has on a gene, the accumulation of genes with respect to both copy number and expression, and the interaction of genes within the system. The programs of the driver gene analysis work in pipeline fashion, where the materialized output from one program is used as input to another later on in the pipeline.

Figure 8 provides an overview of the cancer driver gene analysis. The pipeline starts with the integration of mutation and copy number variation to produce a set of hybrid scores for each sample. The hybrid scores are then combined with protein-protein network interactions to determine effect scores. The effect scores are further combined with gene expression information to determine the connection scores for each sample. The analysis concludes by combining the connection scores across all samples, returning connectivity scores for each gene. The genes with the highest connectivity scores are considered drivers. We now detail each of the steps and conclude with some performance metrics using the two compilation routes.

**Figure 8.**
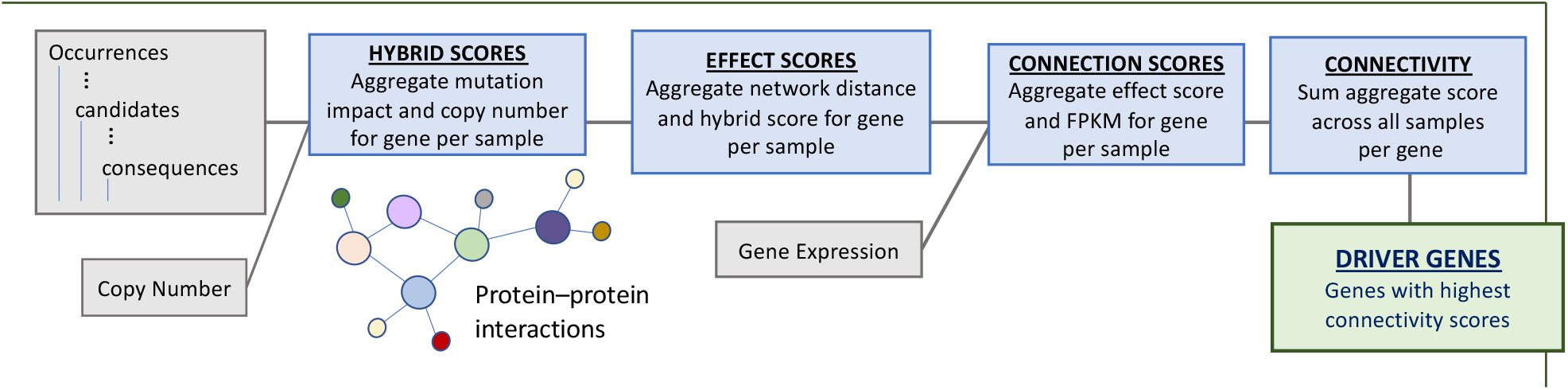
Summary of the cancer driver gene analysis. The pipeline starts by integrating somatic mutations and copy number variation, further integrates network information, and gene expression data. The genes with the highest connectivity scores are taken to be drivers.

#### Hybrid scores

The hybrid score program `HybridScores` is the first step in the pipeline, and is an advanced version of the `SGHybridScores`. The program below describes the process of creating hybrid scores based on the `Occurrences` input. Here, `Samples` provides a map between `sid` and `aliquot` used to join `CopyNumber`, and the hybrid scores are then determined for every `aliquot`. In addition, conditionals are used to assign qualitative scores based on the human-interpretable level of impact (`impact`). The `SOImpact` information is used to integrate values from the nested `consequences` collection into the hybrid score.

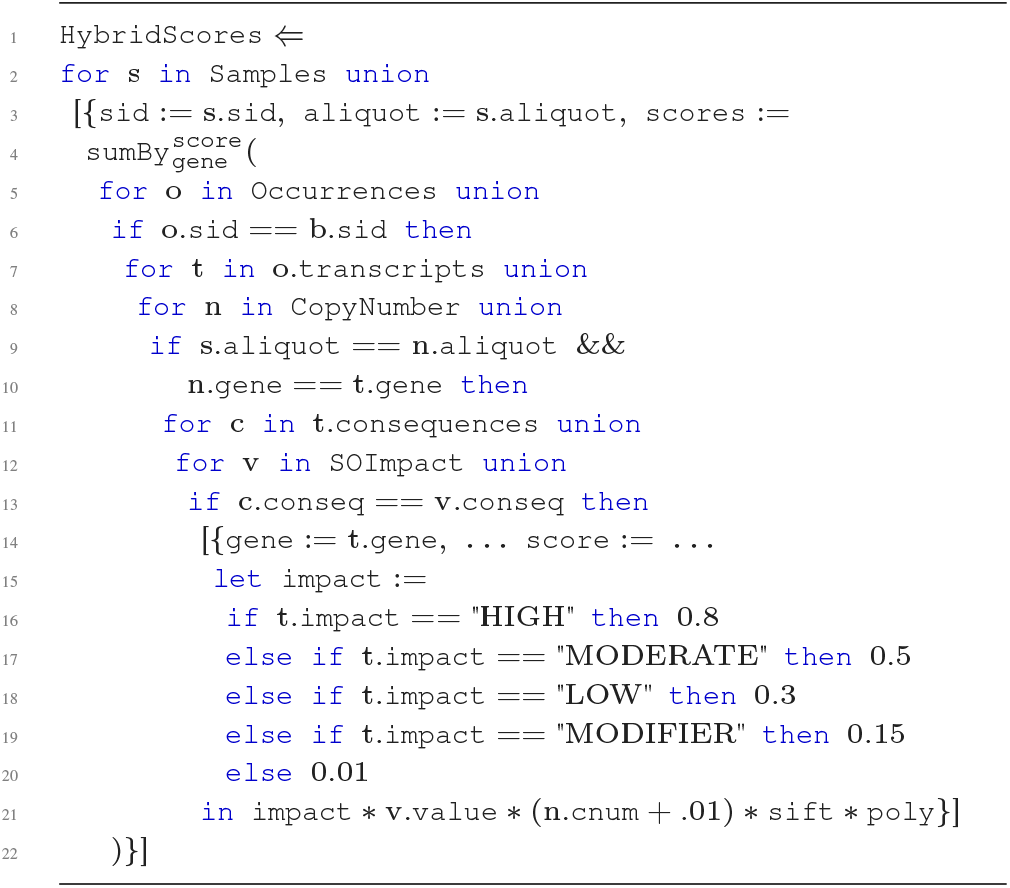

The output type of `HybridScores` is:

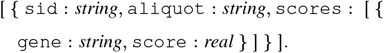

The `HybridScores` program must persist the `aliquot` attribute in order to associate more genomic measurements related to that aliquot later in the pipeline. These hybrid scores now provide a likelihood score of a gene being a driver within a specific aliquot based on both accumulated impact of somatic mutations and copy number variation. The analysis continues to integrate further information to increase the confidence of driver gene scores.

#### By Sample Network

The second step in the pipeline `HybridNetworks` builds individual aggregated networks for each (`sid, aliquot`) pair in the materialized output of `HybridScores`. For each sample, we take the product of the `score` and edge protein `distance` for each edge in the network; genes are associated to proteins based on the mapping provided in the `Biomart` gene map table. The sum aggregate of these values is then taken for each node protein in `Network`, while maintaining top-level sample groups.

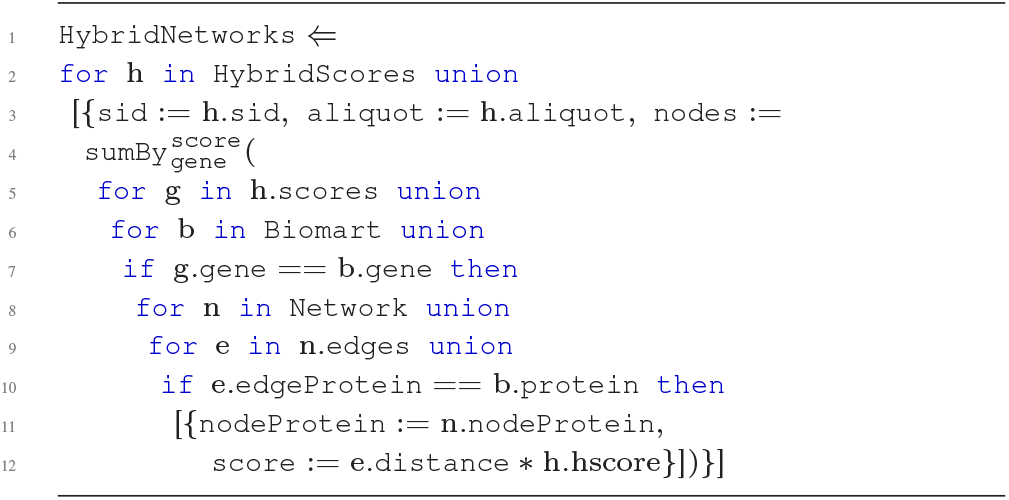

The output type of this query is:

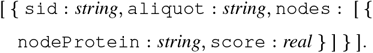

The `HybridNetworks` program produces an intermediate score for each protein in the network by weighting the hybrid scores of nearby proteins in the network (edges) based on their distance scores; thus, this is a partial aggregation of the network data with the hybrid scores using only the edges in the network.

#### Effect scores

To complete the integration of network data with the hybrid scores, the next step is to integrate the nodes in the `Network` to produce effect scores. Effect scores are produced by combining the accumulated edge-based hybrid scores from `HybridNetworks` with the hybrid score for each protein node for each sample in the materialized output of `HybridScores`. As in `HybridNetworks`, genes are associated to proteins using the `Biomart` mapping table.

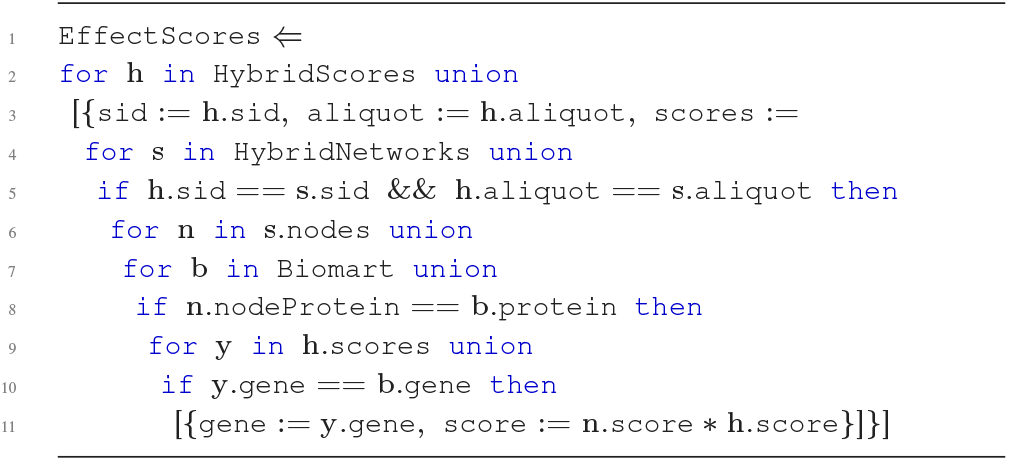

The output type of this query is:

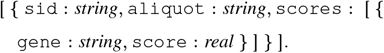

At this point, the effect score is another likelihood measurement for a gene being a driver gene for cancer. The analysis now continues to add confidence to the effect score by further integrating gene-based measurements.

#### Connection scores

The `ConnectScores` program calculates the connection scores. A connection score is the product of the effect score and the FPKM value from the `GeneExpression` table. Gene expression data is combined with the materialized output of `EffectScores` to determine the connection scores for each gene within every sample.

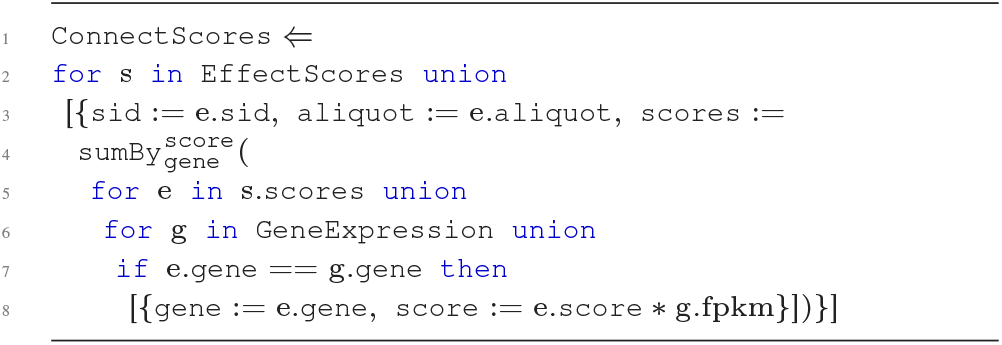

The output type of this query is:

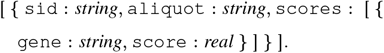

Given the pipeline nature of these queries, the connection scores for each gene are the accumulated somatic mutation, copy number, protein-protein network, and gene expression data for each sample. The connect score can be used to determine the likelihood of a gene being a driver in a specific sample. In theory, this likelihood measurement should have more confidence than the hybrid or effect scores.

#### Gene connectivity

At this point in the analysis, all the genomic measurements have been integrated to produce high-confidence likelihood connection scores for each gene within each sample. The final step is to combine across all samples to identify the highest scoring genes over all samples; this is the gene connectivity. Gene connectivity uses the materialized output of `ConnectScores`, summing up the connection scores for each gene across all samples. The genes with the highest connection scores are taken to be drivers.

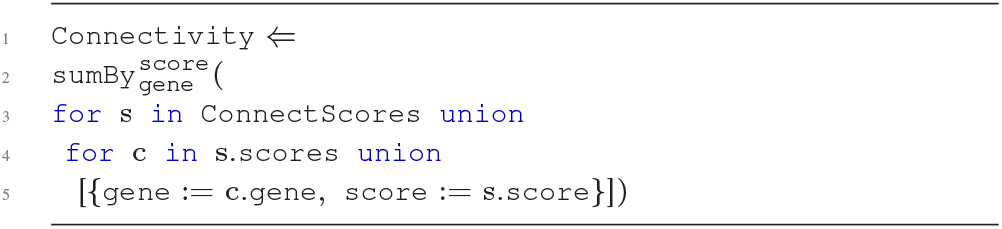

The output type is:

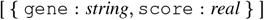

Collectively, these five programs make up the cancer driver gene analysis. The final output of of `Connectivity` is sorted and the top genes are investigated as likely driver genes for cancer. Further confidence can be gained by fine-mapping techniques [28].

#### Runtime performance

The driver gene analysis was executed on a Spark 2.4.2 cluster, Scala 2.12, Hadoop 2.7 on a machine with five workers, each with 20 cores and 320GB memory. We allocate 25 executors per node, 4 cores and 64GB memory per executor, 32GB memory allocated to the driver, and 1000 partitions used for shuffling data. The full TCGA [55] dataset (Pancancer) and the TCGA breast cancer dataset (BRCA) are used. The pancancer dataset uses 280GB of `Occurrences` [24, 33], 4GB of `Network` [50], 23G of `GeneExpression`, and 34GB of `CopyNumber` (34GB). The breast cancer dataset uses the same network information, 6GB of `Occurrences`, 2GB of `GeneExpression`, and 4GB of `CopyNumber`. The runtime of a program is measured by first caching all inputs in memory.

Figure 9 shows the runtimes for the standard (Standard) and shredded (Shred) compilation for the breast and pancancer datasets. Shred exhibits many benefits over Standard. Shred can process the whole of the pipeline on the breast cancer dataset in less than two minutes, a 7x performance gain over Standard. For the pancancer dataset, Shred was able to run to completion for all stages, even when Standard is unable to complete at all. The most expensive stage of the analysis is `HybridNetworks`, which is the combination of the hybrid scores with the network information. This is a nested join that leads to an explosion in the amount of shuffled data. The flattening methods of Standard produces an intermediate join result with 16 billion tuples and shuffles up to 2.1TB before crashing. Shred reduces the size to 10 billion tuples and shuffle to 470GB. These results highlight the benefits of the shredded representation. The shredded representation is essential for scaling up the number samples.

**Figure 9.**
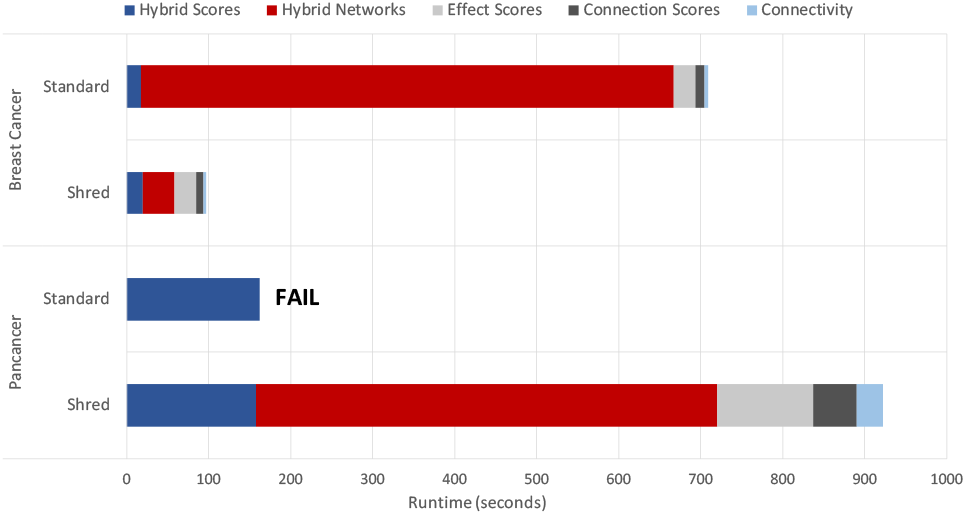
Runtimes for each of the stages of the driver gene analysis. The standard compilation fails atthe `HybridNetworks` program.

Here we have followed the workflow of [60], where the analysis terminates with an identification of driver genes. The three top driver genes reported from our analysis were TP53, FLNA, and CSDE1. All these genes have previously been reported as important for their role in cancer. Future work should explore the pancancer results of this analysis, potentially comparing tumor-site specific driver genes to the identified pancancer driver genes. Naturally, we could also use some of the intermediate scores as features for learning algorithms, as with the previous case study.

### Application 3: Clinical exploratory queries

The identification of personalized diagnosis and treatment options is dependent on insights drawn from large-scale, multi-modal analysis of biomedical datasets. Practical clinical application of such targeted analyses require interfacing with electronic health record (EHR) systems, to provide a data processing environment that supports ease of integrating genomic, clinical, and other biomedical data linked to patients. For example, the Informatics for Integrating Biology and Beside (i2b2) [23] framework facilitates web-based cohort exploration, supporting selection and report generation on clinical attributes. Several proposed solutions for integrating genomic data into i2b2 have been proposed [16, 35, 45]. In these systems, genomic and clinical data are stored in separate databases and then combined in a backend plugin using the i2b2 API.

Figure 10 presents a schematic of an i2b2 instance that supports aggregate analysis with clinical and genomics data sources, i.e. `Occurrences` and `CopyNumber`. The programs of this use case are inspired by such a situation. A user makes a request from a clinical interface. This request represents an analysis that is sent to the backend. The backend application communicates to each of the external data sources to retrieve the necessary data and import them into a Spark processing environment. The application sends the computed results back to the user interface for viewing.

**Figure 10.**
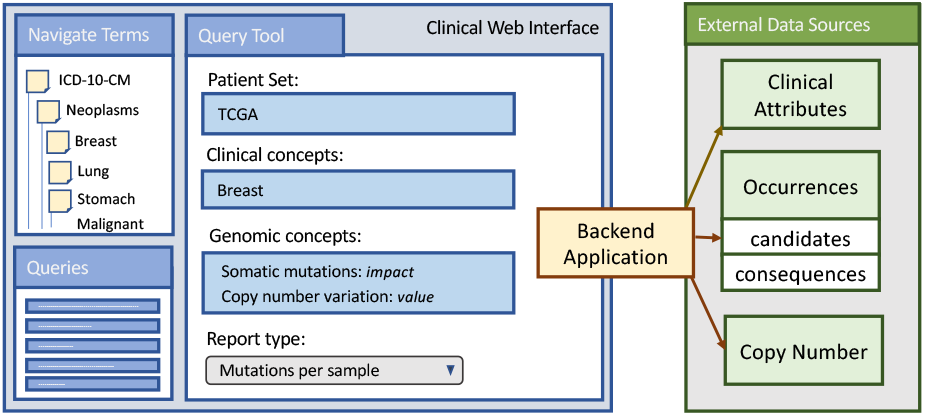
Mock-up of a clinical interface (i2b2) that enables integrative querying of clinical and genomic attributes.

Each of the programs below compromise an analysis that would be performed by the backend application using TraNCE. A major difference from the prior use cases is that here we are not computing just flat aggregates. We are returning nested that will be explored interactively at the web interface. The output will reflect situations where the majority of data fields are returned for exploration by the user,. We now review three such applications, which perform a combination of restructuring, integration, and aggregation of `Occurrences, CopyNumber`, and `Samples`.

### Group occurrences by sample

The `OccurGrouped` program groups the somatic mutation occurrences in `Occurrences` by sample based on `Samples`, producing a collection of nested mutation information for each sample. The program also associates a quantitative value to the consequences at the lowest level in the process, as seen previously in the `HybridScores` program from the driver gene analysis.

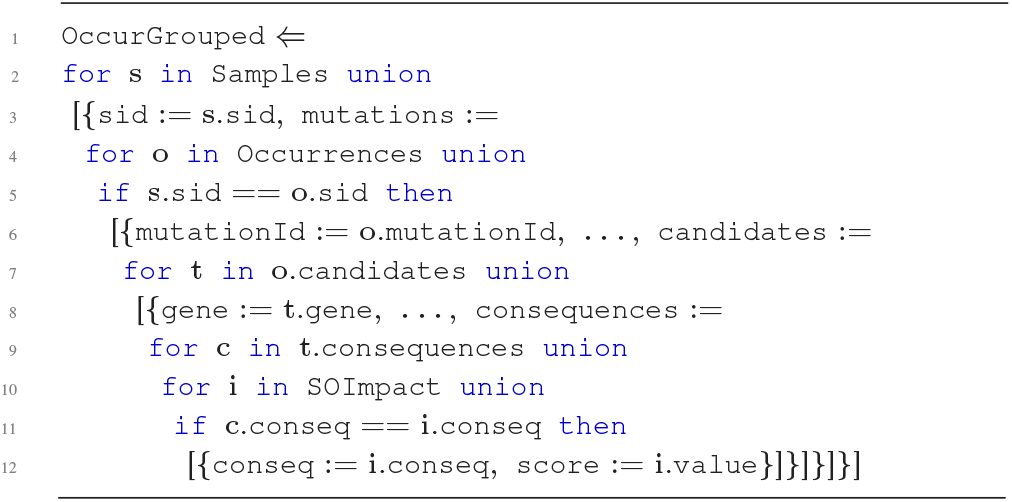

The ellipses represent all the additional fields from `Occurrences`. The output type of `OccurGrouped` is:

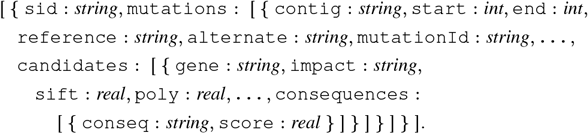

The `OccurGrouped` program groups a mutation data source, like `Occurrences`, based on sample. All information associated to a mutation is returned, with most of the fields of `Occurrences` persisted in the output. The result of this program could feed into a web-interface that provided a detailed view of annotated mutations across a cohort of patients.

### Integrate copy number and occurrences, group by sample

The next program extends `OccurGrouped` by associating copy number information (`CopyNumber`) to each of the genes in the `candidates` collection for each mutation in `Occurrences`. The results are returned group by sample, and the majority of the fields from `Occurrences` are persisted in the output.

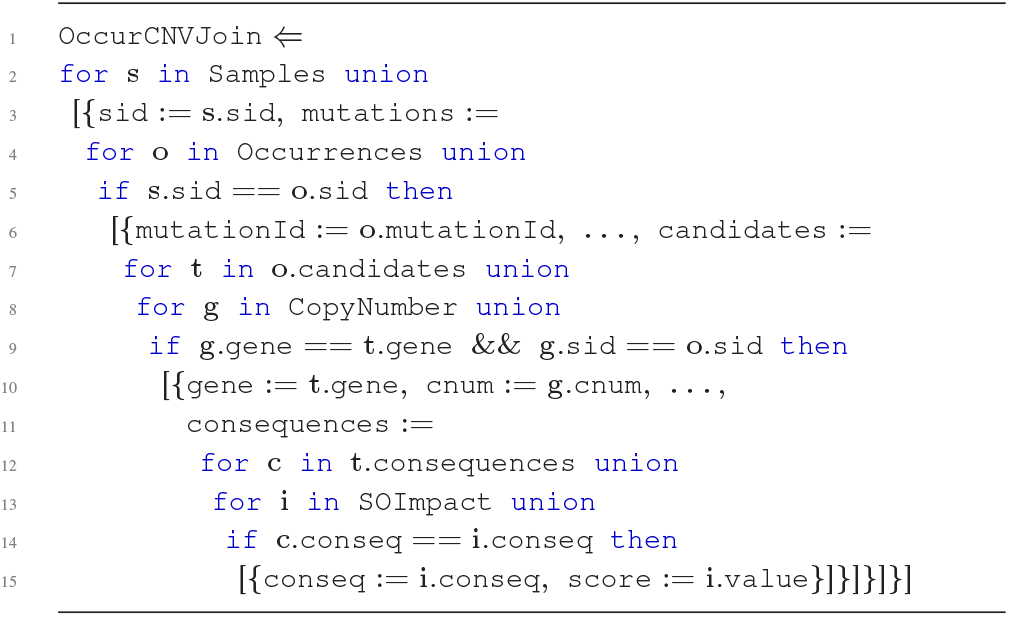

The output type of `OccurCNVJoin` is:

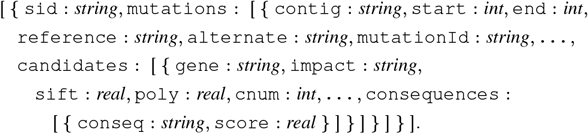

This program exhibits the integration of copy number data on a nested attribute, without any aggregation. The `OccurCNVJoin` program addresses the situation where additional biomedical datasets are integrated for exploration in a consolidated view.

### Aggregate copy number and occurrences, group by sample

The final clinical program combines all aspects of the first two programs and adds an additional aggregation. As in `OccurCNVJoin`, mutations are associated to copy number data to create an aggregate value with mutational impact from the nested `consequences` collection of `Occurrences`. The scores are returned for each candidate gene within each mutation, and the final output is grouped by sample.

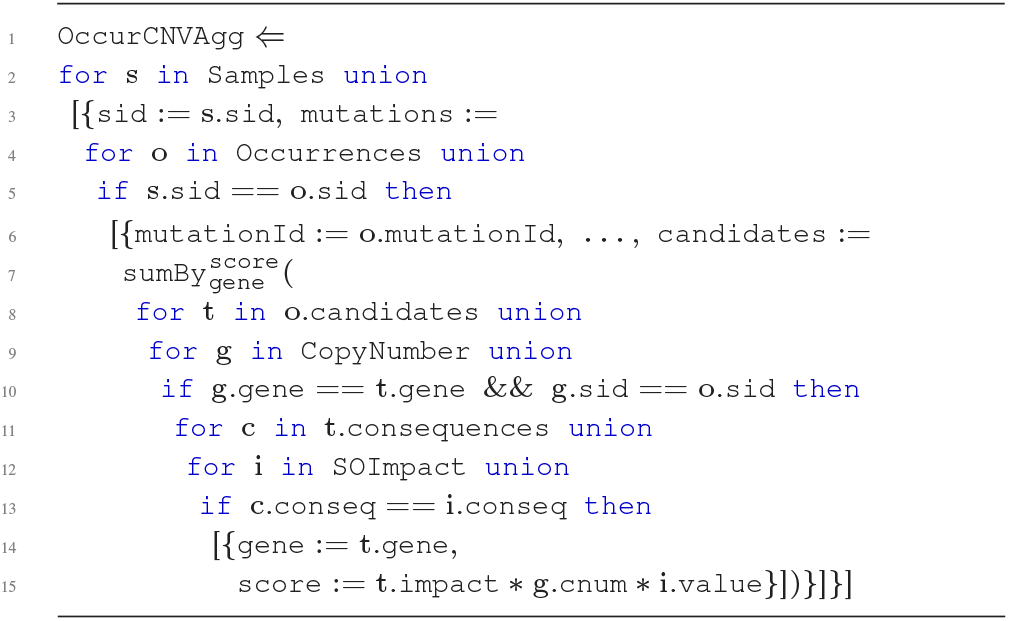

The output type of `OccurCNVAgg` is:

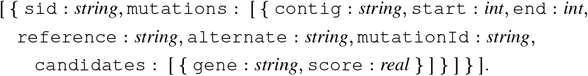

Note that the clinical programs of this use case mimic scenarios that arise from web-based data integration in a clinical setting. Each program adds on a level of complexity - exploring the effects of grouping, joining, and aggregating nested data in a setting that is more exploratory than a research-based analysis. The ability to manipulate these biomedical datasets within a web-based environment that supports frontend clinical exploration presents an interesting application area for the manipulation of nested collections.

### Runtime performance

We execute the clinical programs on a Spark 2.4.2 cluster, Scala 2.12, Hadoop 2.7 on a machine with five workers, each with 20 cores and 320GB memory. We allocate 25 executors per node, 4 cores and 64GB memory per executor, 32GB memory allocated to the driver, and 1000 partitions used for shuffling data. We use the full TCGA [55] dataset (Pancancer) and the TCGA breast cancer dataset (BRCA). The pancancer dataset uses 42GB of `Occurrences` with a 10000 base flanking region, and 34GB of `CopyNumber` (34GB). The breast cancer dataset is 168 Megabytes (MB) of `Occurrences` with 10000 base flanking region and 4GB of `CopyNumber`. The runtime of a program is measured by first caching all inputs in memory.

Figure 11 shows the runtimes for each clinical program, using the standard (Standard) and shredded (Shred) compilation routes. We also include the results for unshredding (Unshred), i.e. the cost of reconstructing the nested output when using the shredded representation. The smaller breast cancer dataset shows performance benefits of the shredded representation (Shred) over flattening methods (Standard). The restructuring in `OccurGrouped` is 80x faster for Shred in comparison to Standard, and still 12x as performant when the nested output type is returned. As operations are added with `OccurCNVJoin` and `OccurCNVAgg`, SHRED exhibits up to 9x performance benefits of Standard. These results show that the shredded compilation method can bring major advantages, even for small-scale datasets.

**Figure 11.**
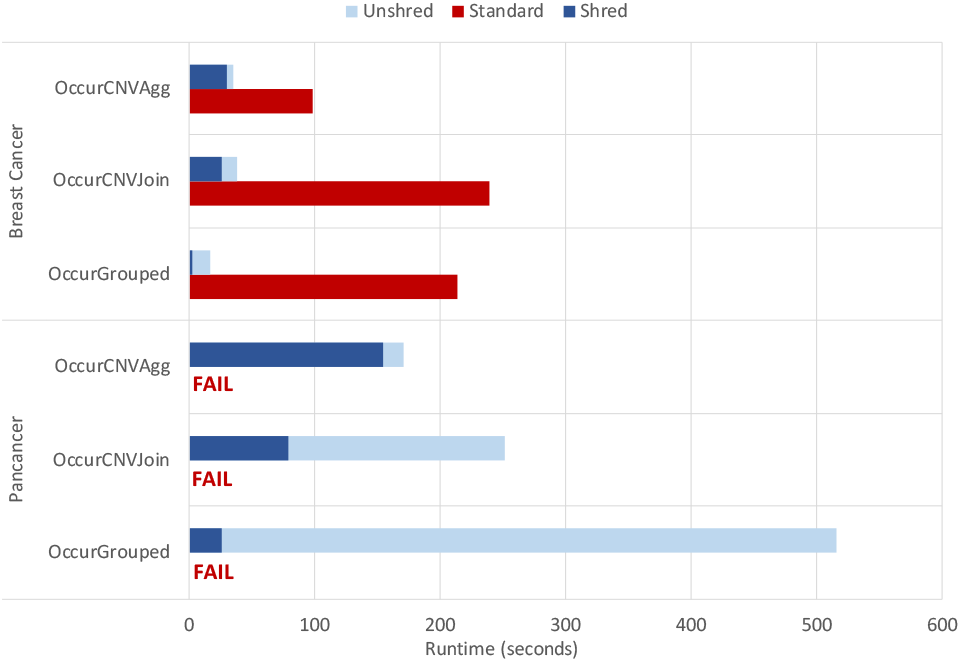
Results for the clinical exploration programs. The standard compilation route fails for all runs with the Pancancer dataset.

For the larger sample set, the results show that Standard is unable to scale, overloading the available memory on the system during each program execution. The results of Shred highlight benefits to the shredded representation. The pancancer `OccurGrouped` is very cheap, but becomes more expensive when the nested output is reconstructed (Unshred); this suggests that the succinct representation used in shredding is essential for scaling. On the other hand, more work is done during the execution of the shredded `OccurCNVAgg` program (Shred), which reduces the cost of unshredding to 3x that of `OccurGrouped`. These results highlight how aggregation in shredded programs can bring further benefits to an analysis even when the output is returned in nested form.

## Sharing in the shredded representation

All the use cases in this section use an `Occurrences` input that is based on the occurrences endpoint of the ICGC data data portal [24], which returns JSON-formatted data following the structure of (3). In this representation, annotations will be repeated within the nested `candidates` collection for mutations that are shared across samples. We can exploit this sharing to create an even more succinct shredded representation of the `Occurrences` data source.

With all somatic mutations are in the `Mutations` data source and all unique annotations in the `Annotations` data source, we can write the following program to construct the data returned from the occurrences endpoint:

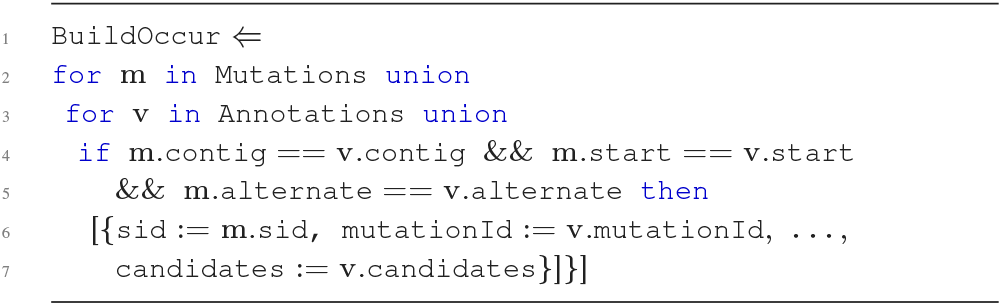

The output type of the `BuildOccur` program matches that of `Occurrences`, presented at(3). The ellipses in the `BuildOccur` program include the additional top-level fields from `Mutations` and `Annotations`. This program describes the construction of the `Occurrences` data source.

### Sharing experiment

To explore the benefits of sharing, we execute the above program using the standard (Standard) and shredded (Shred) compilation routes. We use one somatic mutation (MAF) file from the breast cancer dataset (`Mutations`) containing 120988 tuples, and the associated unique set of 58121 VEP annotations (`Annotations`).

The association in the `BuildOccur` program translates to a join in the compiled Spark application. When the somatic mutations are joined with annotations in STANDARD, the result contains 5170132 tuples nested within the `candidates` collections of the whole output. For Shred, the somatic mutations are joined with the top-level source `Annotations_top`, which has replaced the `candidates` values with labels. The first-level output `BuildOccur_cands` is the same as the input `Annotations_candidates`, which has 3777092 tuples. The shredded representation has reduced the total size of the transcripts by over 1 million tuples.

The results of this experiment are based off a small dataset. Since many of the samples will share mutations specific to cancer, the benefits of sharing will increase for datasets that include more samples. To further explore the benefits of sharing by the shredded compilation route, future experiments should perform the use cases of this section with the output of `BuildOccur` in place of the `Occurrences` data source.

## Conclusions

The TraNCE framework provides a foundation for exploring how query compilation and shredding optimizations can support scalable processing of nested collections. We present several use cases that highlight how the framework can support multi-modal biomedical analyses in research and clinical settings. The results show that the platform has promise in automating the challenges that arise for large-scale distributed processing of nested collections; showing scalable performance for increasing number of genomic variants and performance when flattening methods are unable to perform at all. Further, we exhibit how data integration tasks can feed into machine learning tasks and analytics pipelines. The framework is experimental and its development is ongoing, but our work shows that the techniques applied can provide a basis for many biomedical data integration tasks.

Future work should examine the interface between learning analyses and data integration tasks. For example, a user should be able to describe inference-based tasks within their programs with an extended language that supports iteration and user-defined functions.

The clinical exploration programs present an interesting perspective for the design of biomedical data integration infrastructure. Web-based data types are often nested, and our results show that manipulation of these structures using the standard flattening methods scales poorly. All the use cases have highlighted major advantages for the shredded representation, supporting nested data without compromising the ability to scale. The ability of biomedical systems and analysis applications to work on a succinct representation could present interesting opportunities for optimization, but requires adjustments in backend applications. For example, the clinical exploratory queries could display somatic mutation and copy number data in integrated format to the user, while persisting the shredded representations in the backend. A subsequent request for clinical report generation could use cached inputs that perform localized aggregate operations and return likelihood measurements or risk scores. Future work should consider situations where iterative exploration and aggregation occurs on the data, which is applicable to both research and clinical applications.

Outside of clinical settings, consortium and data biobanks could consider using shredded representations in the backend. Datasets often occur as dump files, which have already gone through a pre-processing phase that employs flattening. Adapting the data representation could support the development of optimized data analysis pipelines. Overall, the TraNCE framework presents an interesting angle for systems development of research and clinical biomedical applications at scale.

## Availability of source code and requirements

- Project name: TraNCE (TRAnslating Nested Collections Efficiently)
- Project home page: github.com/jacmarjorie/trance
- Operating system(s): Platform independent
- Programming language: Scala 2.12
- Other requirements: Spark 2.4.2
- License: MIT Any restrictions to use by non-academics: licence needed

## Availability of supporting data and materials

The data set supporting the results of this article, primarily raw runtimes of performance results, are available in the figshare repository at https://doi.org/10.6084/m9.figshare.13363502.v1.

## Declarations

**List of abbreviations**
API: Application Programming Interface;
BRCA: Breast Cancer;
CNV: Copy Number Variation;
DLBC: Lymphoid Neoplasm Diffuse Large B-cell Lymphoma;
FPKM: Fragments Per Kilobase of transcript per Million mapped read;
EHR: Electronic Health Record;
GB: Gigabyte;
GDC: Genomic Data Commons;
GMB: Gene Mutational Burden;
GTF: General Transfer Format;
i2b2: Informatics for Integrating Biology and Bedside;
ICGC: International Genome Consortium;
ID: Identifier;
JSON: JavaScript Object Notation;
MAF: Mutation Annotation Format;
MB: Megabyte;
MSigDB: The Molecular Signatures Database;
NRC: Nested Relational Calculus;
PMB: Pathway Mutational Burden;
RDD: Resilient Distributed Dataset;
SO: Sequence Ontology;
SQL: Structured Query Language;
TCGA: The Cancer Genome Atlas;
TMB: Tumor Mutational Burden;
TraNCE: Transforming Nested Collections Efficiently;
VCF: Variant Call Format;
VEP: Variant Effect Predictor;

## Consent for publication

Not applicable.

## Author’s Contributions

JS, MB, and MN conceived the idea and design of the framework. JS and MN built the framework. JS and YS conceived and designed the burden based analyses; YS performed and validated the burden-based analyses. JS conceived, designed, performed, and validated the driver and clinical analyses. MB and MN supervised the project. JS wrote the original draft of the manuscript. All authors reviewed and edited the manuscript.

## Acknowledgements

The authors would like to thank Omics Data Automation, Inc. for supplying hardware, compute time, and contributing to use case discussions.

