## Supplementary for "Scalable Analysis of Multi-Modal Biomedical Data"

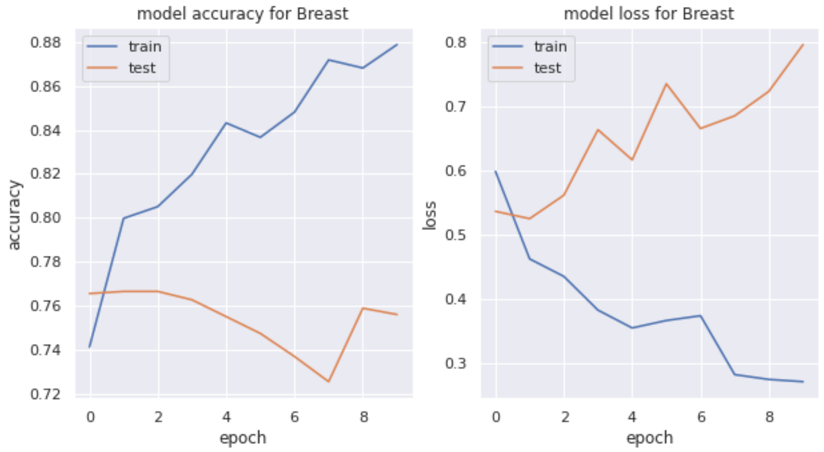


Figure 1: The accuracy and loss of the binary neural network for breast.

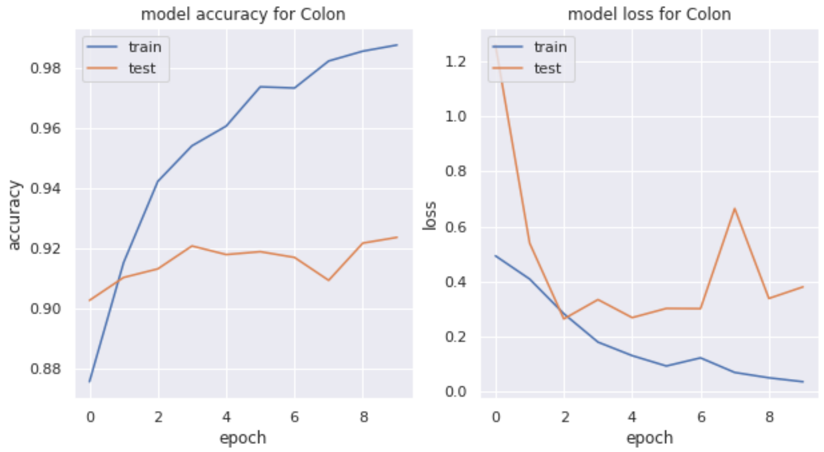


Figure 2: The accuracy and loss of the binary neural network for colon.

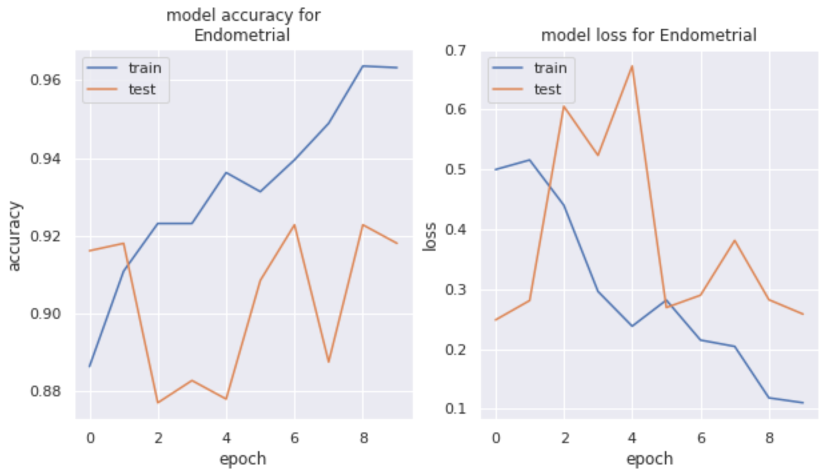


Figure 3: The accuracy and loss of the binary neural network for endometrial.

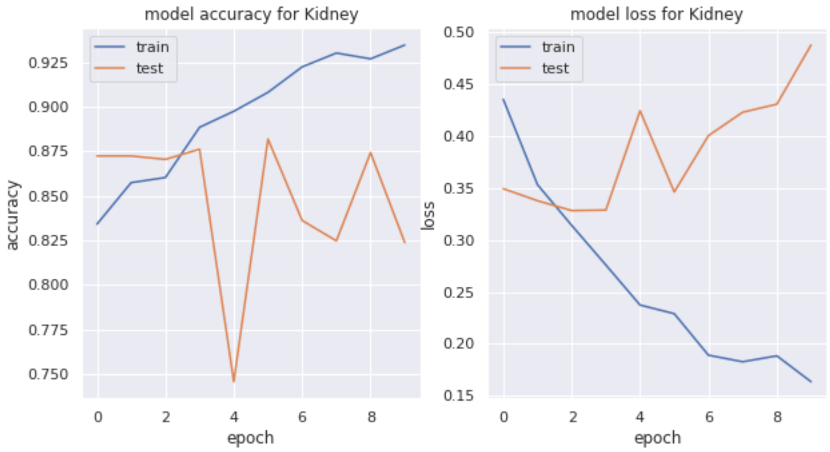


Figure 4: The accuracy and loss of the binary neural network for kidney.

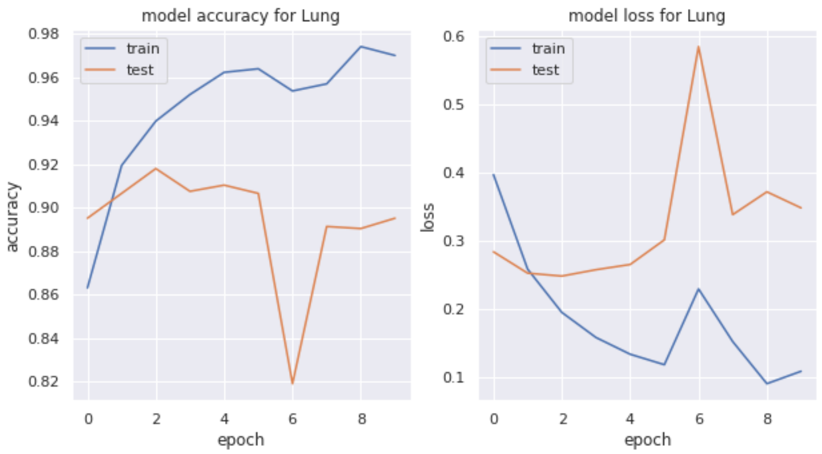


Figure 5: The accuracy and loss of the binary neural network for lung.

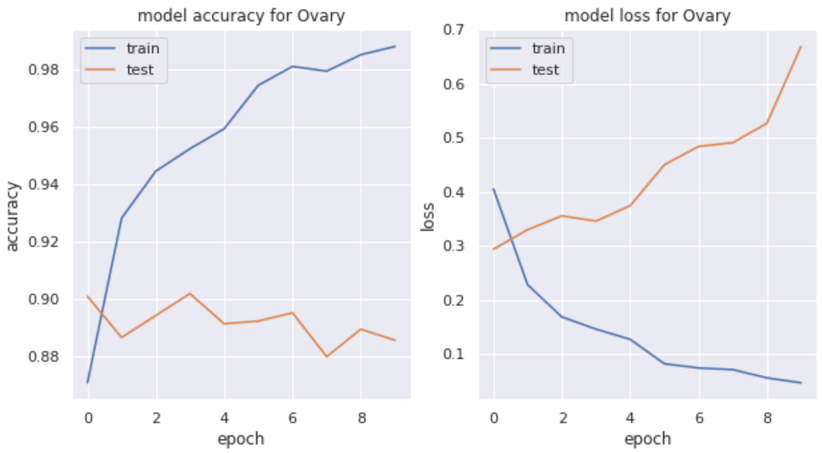


Figure 6: The accuracy and loss of the binary neural network for overy.
